# A 47% decline in wind speed shapes phenology and selection on mismatching in Tree Swallows across a quarter century

**DOI:** 10.64898/2026.10.26.762031

**Authors:** Peter O. Dunn, Linda A. Whittingham

## Abstract

Studies of the effects of climate change on birds typically focus on how temperature affects the average timing of breeding, often neglecting other climate variables such as wind or precipitation and their ultimate effects on reproductive success. Here, we conducted a comprehensive analysis of Tree Swallow (*Tachycineta bicolor*) breeding ecology in southeastern Wisconsin, USA, over a 26 year period (1997– 2023). During this period there was no significant change in mean annual laying date or air temperature. Wind speed was the only weather variable that changed, slowing by 47% during the laying and hatching periods. This decline in wind speed increased food availability (Nematoceran flies), leading to larger clutches, heavier nestlings, and greater fledging success over time. It also allowed later-nesting birds to advance their laying dates. In contrast, earlier-laying birds did not advance their laying date. A changepoint analysis revealed consistently lower food availability and colder temperatures at the end of April than in early May when laying starts. This change in breeding conditions has not shifted during the study and likely acts as a constraint against earlier breeding. Over the years, the distribution of laying dates has been compressed against this early-season “wall”, which has resulted in a dramatic decrease in the variance of laying dates, but relatively little change in the population mean. Slowing wind speed has also relaxed selection against individuals that mistime egg laying relative to peak food abundance (i.e., phenological mismatching). Mismatching exerted relatively weak effects on reproductive success, which was driven mainly by food and weather conditions early in the season, prior to and during laying. This pattern contrasts with other Tree Swallow populations in which reproductive success is primarily influenced by temperature, particularly cold snaps, later in the season during the nestling period. Our study demonstrates that more comprehensive analyses of the effects of climate change are needed because temperature, food and wind speed can interact to produce different effects at the beginning and end of the breeding season, and this can reshape long-term trends in phenology and selection, even before eggs are laid.

## INTRODUCTION

One of the most common responses to climate change is a shift to an earlier start of breeding (Thackeray et al., 2016; Cohen et al., 2018). These phenological shifts have important implications because they could lead to lower reproductive success as a consequence of mismatches between consumers and their food supplies, as well as disruptions in trophic interactions that could impact communities (CaraDonna et al., 2014; Pareja-Bonilla et al., 2025) and ecosystem services (Yin et al., 2023). Although increasing temperatures are leading to shifts in the time of breeding in some species, there are many exceptions (Dunn, 2004; Cohen et al., 2018; Chmura et al., 2019; Buchinger et al., 2022), and understanding why some species are responding, and not others, is important for predicting the impacts of climate change.

There are many possible reasons for a lack of shift (or apparently inadequate shift) in phenology despite warming temperatures, including other conflicting selection pressures (Reed et al., 2013; Day & Kokko, 2015; Visser & Gienapp, 2019), fluctuating selection (de Villemereuil et al., 2020; Le Vaillant et al., 2021), a lack of cues for tracking resource changes (e.g., photoperiod may be the main cue, particularly for migratory species; Both et al., 2010; Romano et al., 2022; Gulotta et al., 2026) and a lack of statistical power (Pujol et al., 2018). Here, we focus on two explanations that have been relatively neglected in the empirical literature. First, it is possible that temperature is not the main driver of phenological changes, although in many cases it appears to be important (79% of species reviewed in Dunn [2004] responded to temperature; Macphie & Phillmore, 2024). For example, precipitation could be a stronger constraint on breeding than temperature, especially in species breeding in tropical or arid environments (Cohen et al. 2018). Similarly, wind speed can be important to species that rely on aerial insects, such as swallows, swifts and bats (i.e., aerial insectivores), because higher wind speed can make food more difficult to catch (Rose, 2009). Among birds, aerial insectivores show some of the largest population declines of any group in North America (Rosenburg et al., 2019), and yet, to date, there have been only a few studies that have examined wind speed in relation to their breeding ecology (Irons et al., 2017). In many cases aerial insectivores are advancing their breeding dates in response to temperature (Taff et al., 2026a), but wind speed could also influence insect abundance independently of temperature. Thus, multiple aspects of weather can affect food availability and reproductive success, but how they operate has rarely been studied in enough detail to understand why some populations respond and others do not.

Another reason for the apparent lack of response to climate change is that populations are changing more subtly through changes in the shape of their phenological distribution, rather than the mean (Vatka et al., 2016; MacPhie et al., 2023). For example, at the population level, variation in breeding time could become smaller if warming conditions allow later breeders to advance, but early breeders are prevented from advancing by poor conditions at the beginning of the season (e.g., cold snaps or low resource availability). Indeed, there is increasing evidence that breeding seasons are becoming shorter in birds (Dunn & Møller, 2014; Hallifors et al., 2020). Although there is much discussion of an ecological or physiological “wall” early in the season (Winkler et al., 2002), it has rarely been explicitly tested or examined in terms of its ultimate effect on reproductive success or its long-term consequences for the response to climate change. If early season constraints are strong enough, then the population mean time of breeding may not change, even though a significant part of the population (in the right tail of the distribution) is advancing. In other words, the distribution of breeding can be compressed and have a stronger peak if the climate is changing and there are constraints on breeding earlier.

This inability to shift the distribution earlier may have consequences for reproductive success. In particular, the mismatch hypothesis suggests that if populations do not shift their time of breeding to match the availability of resources they need for breeding (e.g., food to supply growing offspring), then the mismatch in timing will lead to decreasing reproductive success. Currently, there is mixed evidence to support the mismatch hypothesis (Dunn et al., 2011; Renner & Zohner, 2018; Dunn, 2019; Samplonius et al., 2021; Zhemchuzhnikov et al., 2021; Kharouba & Wolkovich, 2023; Youngflesh et al., 2023). Although these studies used different types of data and criteria, they often agree that we need to test our assumptions better, including the assumption that we are measuring the weather and food variables that are most relevant to the reproductive success of a particular population.

Here we present a comprehensive analysis of the interacting effects of weather and food availability on the phenology and reproductive success of an aerial insectivore, the Tree Swallow (*Tachycineta bicolor*). Numerous studies have shown that the breeding ecology of Tree Swallows is closely linked to flying insect availability or weather (e.g., Nooker et al., 2005; Twining et al., 2018; Berzins et al., 2021; Garrett et al., 2022a). However, our long-term study revealed that the distribution of egg-laying has been squeezed by changes in weather and food availability into a narrower distribution with important reproductive consequences that would not have been fully understood by simply studying the mean annual response to temperature. Crucially, a traditional analysis focused on mean timing and temperature could have led us to the incorrect conclusion that our population was not responding to climate change.

Like many aerial insectivores, Tree Swallows are declining overall, but the decline is primarily in the northeastern part of their range (Nebel et al., 2010). Our study population in the midwestern United States has been relatively stable in terms of the number of breeding pairs (see below; Shutler et al., 2012). Across their range, Tree Swallows are also advancing their laying date in response to increasing temperature (Dunn & Winkler, 1999), but the laying date response is variable across their range because, in part, some populations are experiencing more warming than others (Taff et al. 2026a). Interestingly, laying date in different populations has a similar response to temperature (i.e., their sensitivity is similar; Taff et al., 2026a), but some populations appear to be more sensitive to wind speed than temperature, at least in terms of laying date (Irons et al., 2017). Furthermore, later in the season, nestling growth and survival are related to food abundance in some populations (McCarty & Winkler,1999; Twining et al., 2018), but not others (Dunn & Roberston, 1992; Imlay et al., 2017). Thus, our goal was to examine the entire breeding cycle from laying to fledging in relation to food availability and weather (temperature, wind, rain), because it is not clear how and when different environmental variables affect the reproductive success of swallows and, thus, the intensity of selection on timing of breeding. Understanding why populations have different responses to these environmental variables will be important for predicting their response to climate change and why some populations change while others do not.

## METHODS

### Study species and area

Tree Swallows (18-22 g) are a common breeding species in the northern United States and Canada. They typically use secondary cavities in trees as nest-sites, but they will also use nest-boxes, which has made them a widely studied species. At our study site in southeastern Wisconsin, USA, swallows arrive on the breeding grounds in late March to mid-April, several weeks before most egg-laying starts in mid-May. This early arrival is likely due to competition for nest-sites which are limited in number (Stutchbury & Robertson, 1987). There is usually a single brood each season, and clutches of 5 and 6 eggs are most common (normal range = 3-8; Robertson et al. 1992). Incubation starts on the day the penultimate egg is laid and lasts for 12-15 days. After hatching, both parents feed the young, which remain in the nest for 18-22 days (Robertson et al., 1992).

We studied Tree Swallows from 1997 – 2023 in a box-nesting population at the Field Station of the University of Wisconsin-Milwaukee in Saukville, Wisconsin, USA (43° 23’N, 88° 01’W). To facilitate comparisons between populations, the boxes in our grids were the same dimensions and were arranged in a similar grid pattern (28 m nearest neighbor distance) as boxes in a long-term study at the Queen’s University Biological Station in Ontario, Canada (44° 34’ N, 76° 20’ W; Robertson C Rendell, 1990). Our grids of boxes were set up in two hayfields 0.7 km apart. Excluding the first year, the number of available nest-boxes varied from 71–90 each year (average = 83) and, on average, 95% were occupied each year (range: 77-100%).

### Nest box monitoring

Nest boxes were checked at least every other day from the day nesting material reached 4 cm until clutch completion, and again from day 13 of incubation until the nestlings were 12 days old (“day 12”). Nestlings were weighed and measured on day 12, which is the age when nestlings typically reach their asymptotic body mass before fledging (Zach & Mayoh 1982). These observations enabled us to determine laying date, clutch size, hatch date, and the weight and number of young on day 12. Fledging success was estimated as the number of young in the nest at 12 days of age minus any dead young found in the nest after fledging (usually checked at 18-20 days).

Female Tree Swallows usually lay one egg per day, although interruptions of 2-3 days can be common during rainy or cold weather. Longer interruptions are usually due to abandonment and takeover by a new female that covers the previous eggs with new nesting material and then starts laying again. Here we focus on first nesting attempts and avoid these replacement clutches by excluding any clutch that showed evidence of takeover by a new female including interruptions during laying totaling more than five days, unusual egg-laying (2 eggs in one day) or new females (based on banding). We also excluded any nests in which experiments might have influenced laying date or other reproductive parameters (see Results for sample sizes).

Adults were caught inside nest boxes, measured (wing chord and tarsus length), weighed and marked with a numbered aluminum U.S. Geological Survey band on the right leg and a colored plastic band on the left leg. Body condition was estimated from the residuals of the regression of body mass on wing length and capture date (to account for seasonal changes). Females were classified as second-year (SY) or after second-year (ASY) on the basis of plumage coloration Hussell (1983).

### Insect sampling and food availability

We sampled insect abundance with a tow-net identical to Hussell and Quinney (1985) during the first three years of study (1997-1999) and a 1.5 m tall suction trap (fig. 4.2 in Southwood C Henderson 2000) for the remainder of the study (2000-2021). Except for the date of peak insect abundance, we exclude the three years of tow-net sampling so the estimates are comparable; this resulted in 15 years of samples from the suction trap. Each day traps were operated for at least 80% of daylight hours. Daily sampling started in late April or early May (mean start date = 24 April; range of first dates = 9 Apr to 7 May) and finished in late June (mean end date = 27 June; range of end dates = 22 June to 2 July). Sampling with the suction trap occurred on an average of 65 days per season, and there were 956 days of sampling across all 15 years.

To estimate insect biomass, we counted the number of insects in different taxon and size categories, and then multiplied that number by its average dry mass (Hussell C Quinney, 1982; Dunn & Hannon, 1992). Our taxa for analysis included flies in the suborder Nematocera (primarily midges, mosquitoes and craneflies), other Dipteran flies (primarily Brachycera), Lepidoptera, Coleoptera, Hymenoptera, Homoptera and Ephemerata. In this study we focus primarily on Nematoceran flies because these contain high levels of polyunsaturated fatty acids (PUFA) that provide high quality food for growing Tree Swallow nestlings (Twining et al., 2016) and their abundance is related to fledging success (Twining et al., 2018).

There were 20 missing days of insect sampling in the first 4-6 days in May during 2000-2003, so we imputed values for Nematocera biomass using predicted values from a regression equation (Appendix S1). Excluding these imputed values did not have a qualitative effect on the results. We excluded the last four days of insect sampling in 2008 (June 23-26), because they included outliers for Nematocera biomass (85 – 779 mg) that were above the 99.5th percentile (55.4 mg). Most nests had already fledged young by this time of year, so excluding these days had no qualitative effect on our measures of reproductive success. Prior to analysis, insect abundance was Ln+0.001 transformed. The peak date of insect abundance was identified as the maximum of a 5 day running average and visually verified by plotting each year and fitting the data with splines.

### Weather data

We used an automated weather station located halfway between our two grids of nest-boxes to collect most of the weather data. Some weather data were also obtained from the Milwaukee International Airport (61 km away) for comparative purposes and to fill in missing data from the UWM Field Station (see Results). Weather variables included daily air temperature (mean, maximum and minimum), wind speed (m/s), rainfall (mm), and solar radiation (Kw/m^2^).

### Statistical analyses

Most analyses were conducted in JMP v. 19 (JMP® 19, 2026), with some additional specialized packages in the R 4.4.2 programming language (R Core Team, 2024) as described below. For purposes of reproducibility, most analyses were redone in R and all of the R code and data files have been deposited in Zenodo (see Data Availability Statement). Google AI mode was used to troubleshoot R coding, but all code and results were verified manually.

#### Stages of the breeding cycle

To examine long-term changes in weather and food abundance on swallow breeding parameters, we initially focused on changes in weather or food on or around (5 or 10 days before) the long-term average date for laying or hatching (see below for justification). For some analyses we also used four biologically relevant time windows (stages) that were informative in previous studies: the pre-laying, laying, incubation and nestling periods.

The pre-laying period was defined as 1-15 May to examine the environmental factors influencing egg formation and the decision to start laying (laying date). We chose 1-15 May based on strong correlations between laying date and temperature during this period in several populations (Hussell, 2003; Dunn et al., 2011) and for consistency with a similar study of the effects of both temperature and insect abundance at Port Rowan, Ontario (LeClair, 2012). We also examined three other 5 day windows (1-5, 6-10, 11-15 May), and based on AICc values, the 1-15 May period provided the best window for predicting laying date using insect (Nematoceran) biomass (see Results).

In contrast to laying date, there was less a priori information about the weather variables associated with the peak date of insect biomass, so to find the best predictor of the peak date, we used a sliding window approach in the R package climwin (Bailey C van de Pol, 2016). Note that climwin is not designed for variables that are unrelated to a date, such as clutch size, fledging success or insect biomass. For these other variables, we had to use time windows based on breeding activity such as the “laying” period, rather than a single date.

We defined the laying period for each population and year as the 25^th^ to 75^th^ percentiles of laying dates (see also Dunn et al., 2011). The 25^th^ percentile avoids extremely early nests and represents the beginning of laying better than the mean laying date. Food abundance during the laying period was estimated for each population and year using the average insect biomass (log-transformed) during the five days prior to the 25^th^ percentile of laying dates to the 75^th^ percentile of laying dates. We used a five day period before the 25^th^ percentile because rapid yolk synthesis averages five to six days in Tree Swallows (Arida et al. 2006), and, thus, five days allowed us to capture the effects of food on laying throughout the period of rapid yolk accumulation and the formation of the remainder of the egg (see also Nooker et al., 2005; Ardia et al., 2006). We also examined a wider window for the laying period (2 days before the 10^th^ percentile to the 90^th^ percentile) that was used by LeClair (2012) at Long Point, Ontario, but the results were qualitatively similar.

The incubation period was defined as the period from the 50^th^ percentile of clutch completion to the 50^th^ percentile of hatch dates, consistent with LeClair (2012). Lastly, the nestling period was defined as the period from the 25th percentile of hatch dates to 15 days after the 75th percentile of hatch dates (Dunn et al., 2011). We used 15 days after hatch to define the end of the nestling period as fledging can occur after this age and nestlings have reached a plateau in body mass (Zach & Mayoh, 1982; Winkler & Adler, 1996).

Our initial analyses of laying date and temperature revealed no long-term change when we examined annual means; however, there was evidence that later nesting birds (>75^th^ percentile) were advancing their laying dates and this was changing the shape of the laying date distribution. We also found that there has been a dramatic (47%) decline in wind speed at our study site, which has increased food availability for swallows. To examine this in more detail, we switched to analyses that could examine both the mean and shape of the egg-laying distribution.

#### Analysis of laying date using Bayesian distributional mixed models

We used Bayesian distributional mixed models in the R package brms (Bürkner, 2017) to evaluate long-term shifts in the mean and shape of the laying date distributions as well as constraints on early breeding and their consequences for reproductive success. Using mixed models has several advantages over the analysis of annual means above (*n* = 25 years), because we gain increased statistical power from using all of the individual nesting observations (*n* = 1549 nests), and the models can account for individual variation by incorporating a random effect for each female (using “ 1| female_id” in the model). For the initial models examining temporal changes, laying date was modelled using the Gamma distribution which allowed us to examine both the mean and shape of the distribution in terms of variance and skew. The model used a log link function, so the regression coefficients ( *β*) represent exponential trends that were transformed to percentage change using: % Change = (e *^β^*-1)*100. We transformed our Julian laying dates by subtracting 123, which meant that our earliest recorded laying date, 4 May (day 1), was the start of the season. This ensured that percentage change was calculated relative to a biologically relevant starting point rather than an arbitrary calendar date (e.g., January 1).

Next, we examined environmental variables that might influence the mean and shape of the laying date distribution over time. Here we began with a Gamma distribution model for laying date that contained year, weather (temperature, wind speed and rain) and food (nematoceran biomass) as predictors and the random effect of individual female (female id). Based on the decline in wind speed that we found in our preliminary analyses, we also included two interactions: 1) pre-laying wind speed and food abundance and 2) pre-laying wind speed and year. This allowed us to test the effect of wind speed on food abundance and changes over time. All predictor variables were standardized before analysis. In this initial full model we had two types of weather and food variables. Our absolute variables were averaged for the pre-laying period (1-15 May) and, thus, were the same for all females within a year, but different between years, while our relative variables were averaged for the 5 days before each female started laying. This allowed us to examine variables influencing both the population level changes (absolute) and shorter-term individual responses (relative). Relative variables are not as useful for explaining changes in mean and skew, but they can be useful if birds are responding to environmental conditions just before laying (5 days in this case), instead of earlier in May during the pre-laying period, which can potentially be important for birds starting to lay later in May. To test the influence of certain types of variables, such as those affecting the shape of the distribution versus the mean, we ran different models and compared them with a LOO (Leave-One-Out) procedure (loo package, Vehtari et al., 2017) and the Watanabe Akaike Information Criterion (WAIC) which estimates a model’s predictive performance similar to AIC (lower values indicate a better model). In comparisons of models, brms provides the difference in Expected Log Predictive Density (elpd_diff) between two models and whether that difference is statistically reliable based on its standard error (se_diff). We also estimated a Bayesian R^2^ in the rstantools package (Gelman et al., 2019).

For each model, we initialized four independent parallel Markov chains. Each chain was run for a total of 3,000 iterations. The first 1,500 iterations of each chain were discarded as a warm-up period to allow the sampler to locate the target posterior space and optimize the step size (epsilon) and mass matrix. A total of 1,500 post-warmup draws were retained per chain, yielding a combined total pool of 6,000 posterior samples for final inference. To improve sampling in some analyses, we slowed the step size by setting the argument “adapt_delta” to 0.9999 and increased the tree depth to be evaluated by setting “max treedepth” to 15. Posterior distributions were estimated using the No-U-Turn Sampler (NUTS), an extension of Hamiltonian Monte Carlo (HMC) sampling (Bürkner, 2017). Chain convergence was verified by R-hat values equal to 1.00 and large effective sample sizes (ESS typically > 1000). We considered parameters significant when the Bayesian 95% credible interval (CrI) did not overlap with zero.

#### Change point analyses

To determine if there were constraints on the timing of laying early in the season, we looked for significant changes in temperature and food availability using changepoint analysis in the multiple change points (mcp) package in R (Lindeløv, 2020). We assumed that significant shifts in the intercepts of regressions of temperature or food on date would be indicative of environmental thresholds that could restrict breeding earlier in the season. We chose mcp because it allowed us to find changepoints along a day of the year timeline while incorporating year-to-year shifts by adding year as a random effect in the model. The mcp package does not explicitly test for the number of changepoints, so we compared models with 0, 1 and 2 changepoints using leave-one-out cross-validation and the Expected Log Predictive Density Difference (elpd) in the loo package (Vehtari et al., 2017). The top model receives a elpd value of 0 and all other models receive a negative number; if the number is less than –4, then the top model is considered significantly better.

#### Testing for temporal shifts in the constraints on early breeding

To determine whether the earliest edge of the laying date distribution has been shifting across years (“flexible wall”) or has a relatively fixed boundary (“hard wall”), we examined the 5^th^ quantile of laying dates as an index of the start of laying using the quantreg package in R (Koenker, 2025). Our full model was similar to the Bayesian model above for the analysis of the mean and shape of the full laying date distribution, but we only included the absolute prelaying weather and food variables. In particular, we estimated the response of the 5^th^ percentile of laying dates to standardized (“scaled”) predictors of time (year_scaled), pre-laying weather (preLayTemp_scaled, preLayRain_scaled, preLayWind_scaled), food abundance (preLayNemato_scaled), and the two-way interaction terms with wind (preLayWind_scaled × preLayNemato_scaled and year_scaled × preLayWind_scaled) that were modeled previously. To account for the non-independence of repeated measures across years for the same individuals, we estimated clustered bootstrap standard errors and *p* values across 2,000 bootstrap replications of individual females (female_id). A flexible boundary wall for the 5^th^ percentile of laying date was tested by evaluating the significance of time (year_scaled) and its interaction with pre-laying wind speed (year_scaled:preLayWind_scaled); in this case a statistically non-significant temporal response would indicate a fixed threshold across the study period.

#### Selection

To examine selection on laying date, as well as the mismatch between hatching date and peak food abundance, we calculated linear and quadratic selection gradients following the regression approach of Lande and Arnold (1983). Linear selection gradients tested whether selection favors all birds to breed earlier in terms of absolute calendar date, while quadratic selection gradients tested whether there is stabilizing selection around an optimum, such as the date of peak of food abundance. We used relative fledgling success (individual fledging success divided by the mean success for that year) as our measure of relative annual fitness because natal recruitment rates are low (<5%) in our populations. For these analyses, traits (laying date or mismatch) were standardized across all years by subtracting each observation from the mean of the trait and dividing by the standard deviation (mean = 0, SD = 1). The model included relative fledging success as the response and the fixed effects of the trait (laying date or mismatch) as both a linear and quadratic (trait squared) term, a continuous time variable (decade = year/10), female age (ASY, SY or unknown) and the interaction between the linear and quadratic terms and decade. Decade was used as a continuous time variable to test for long-term trends within the same model that was testing for selection, while year was included as a random factor to account for unmeasured annual variance. We incorporated both random slopes and intercepts for the trait variable across years (i.e., (1+Trait∣Year)) to account for annual fluctuations in directional selection, which helps to ensure that the longer term trend was not driven by a few extreme seasons. Selection gradients were estimated as the slope of the regression of relative fledging success on standardized laying date or mismatch. To obtain stabilizing selection gradients similar to those of Lande and Arnold (1983), we multiplied the partial regression coefficients of all quadratic terms by two (Stinchcombe et al., 2008). Similar to the Bayesian analyses above, we implemented the models using the brms package, but with a Gaussian error distribution. For the long-term temporal trends in selection, we also calculated the probability of direction (pd), which measures the proportion of the posterior distribution that has the same sign as the median (Makowski et al., 2019). In our case, it gives the probability that directional or stabilizing selection has changed over time, ranging from 0.50 (complete uncertainty) to 1.00 (absolute certainty). Due to the high inter-annual variance in long-term datasets, 95% Bayesian credible intervals can occasionally overlap zero even when there are strong continuous trends (de Villemereuil et al., 2020). Thus, we used probability of direction (pd) as a continuous metric of statistical evidence, as it explicitly quantifies the posterior certainty of directional and stabilizing selection without relying on arbitrary significance thresholds.

#### Structural equation modeling of reproductive success

We were interested in how food and weather affect each stage of reproduction from laying date and clutch size up to the number of young that fledge. Due to the many possible ways in which environmental variables could affect reproduction, we used a structural equation model (SEM) in JMP to determine the relative importance of food and weather at various stages of reproduction. We constructed the SEM based on previous results and a priori hypotheses about the relationships between each environmental variable and reproduction (see Supplement for hypotheses and their justification). To simplify the analysis, we only used known (banded) adult (ASY/AHY) females with data on all reproductive parameters (n = 548 nests). We started with an SEM (Appendix S1) that contained food (Nematocera biomass), temperature and wind variables that were likely to affect reproduction, including laying date, clutch size, number of eggs hatched (N hatch), number of young reaching 12 d of age (D12) and number of young fledged (N Fledge). We averaged food, temperature and wind speed for the four reproductive periods defined above: pre-laying, laying, incubation and the nestling period. All variables were standardized before analysis.

## RESULTS

### Is Laying date changing with year?

We examined 1549 first nesting attempts over a 26 year span (1997 – 2023). This sample did not include nests with experiments or changes in occupants that might have biased reproductive parameters. We did not collect nesting data in 2007, so there was a total of 25 years of data (mean ± SE = 62 ± 3.4 nests per year; interquartile range = 61 – 79 nests). On average, females laid the first egg of their clutch on 18 May (SE = 0.13 d, interquartile range = 14 – 21 May; range = 4 May – 11June. Figure 1), which was 17 days before the peak of biomass (using a running 5 day average) for nematocerans (4 Jun, SE = 3.7 d, interquartile range = 22 May – 21 June) and 28 days before the peak for all insects (15 Jun, SE = 4.2 d, interquartile range = 13 – 26 June; Figure 1 top panel).

**FIGURE 1.**
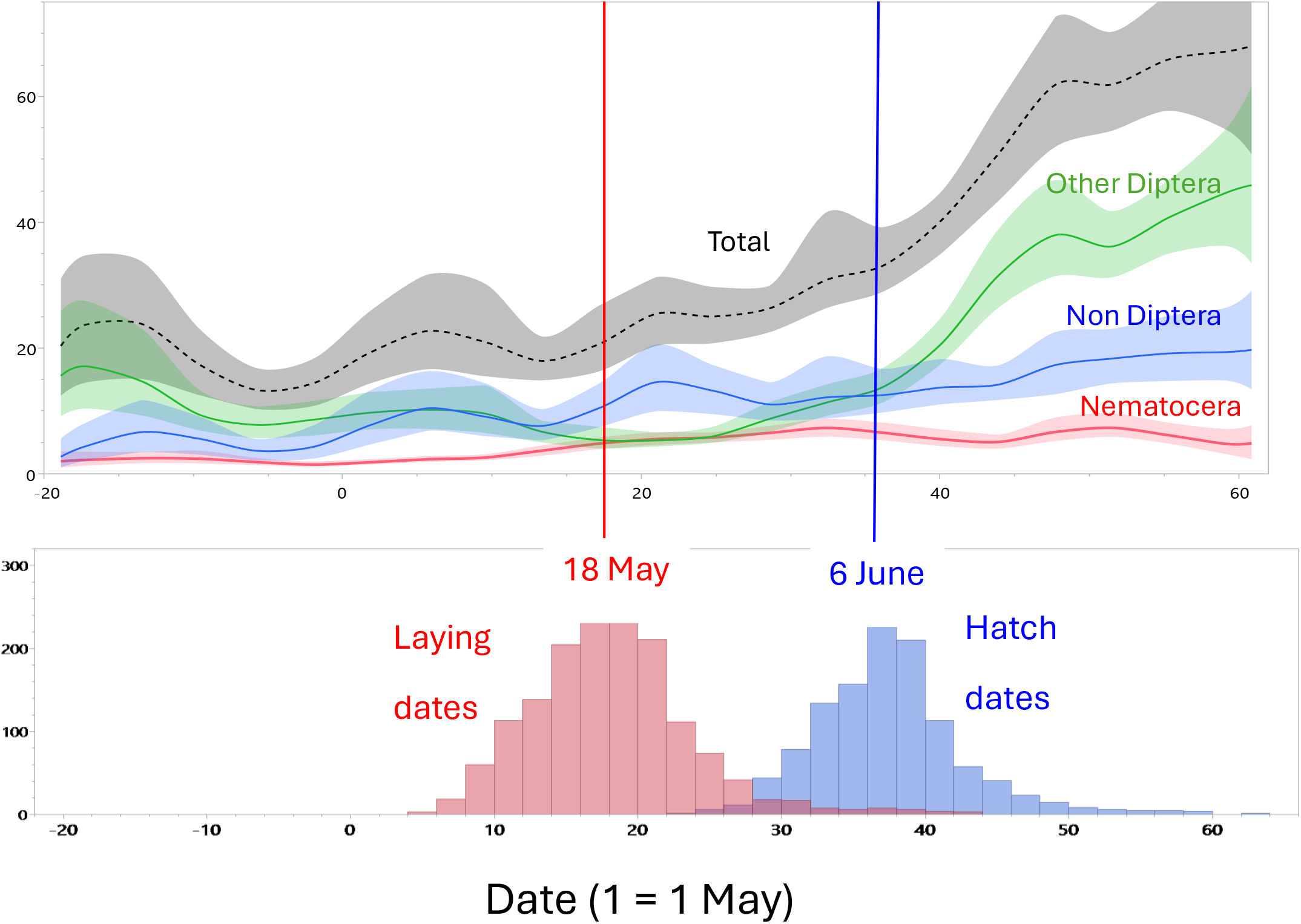
Insect biomass (mg/day) (top panel) and the distribution of egg laying (red bars) and hatch (blue bars) dates (bottom panel) during spring (1 = 1May) in Saukville, WI, USA (2000-2021). Biomass data were collected with a suction trap on 956 days during 15 years. Data from 1997-1999 are not shown here because insect biomass was collected with a different type of trap (tow-net).

Laying date has advanced significantly by 1.4 days during 1997-2023 when we analyzed individual nests in a simple linear regression (Figure 2A; *n* = 1549 nests; slope = – 0.057 d/yr, *t* = 2.99, *p =* 0.003). However, when we analyzed mean annual laying date over the 26 years of study (ie. n= 25 years of data), the advancement (3.7 d or 19%) was not significant (Figure 2B; –0.144 ± 0.08 d/yr; *r*^2^ = 0.10, t = 1.64, *p =* 0.114). Annual mean air temperature has also not changed over time (Figure 2B).

**FIGURE 2.**
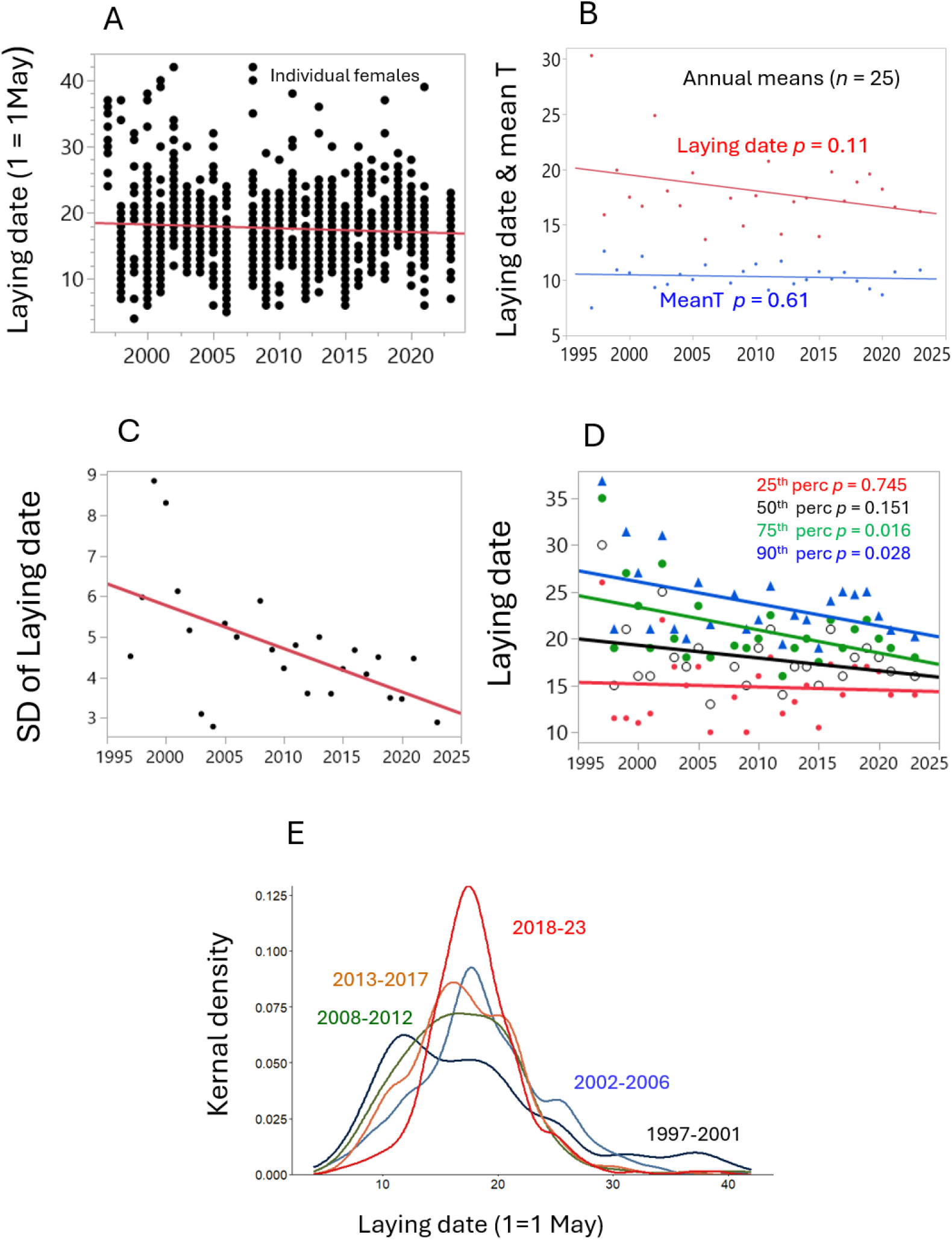
Laying dates of individual females (n = 1549 nests) **(A)** have advanced over time (slope = –0.057 d/yr, *t* = 2.99, *p =* 0.003), but there has been no change in the annual mean laying date (–0.144 ± 0.08 d/yr; *r*^2^ = 0.10, *t* = 1.64, *p =* 0.114) or mean air temperature (MeanT; –0.065 ± 0.05 d/yr; *r*^2^ = 0.07, *t* = 1.30, *p =* 0.206) **(B)** across the 25 years of study (1997-2023). No data were collected in 2007 or 2022. **(C)** Variation (SD) of laying date declined across the study period (–0.108 ± 0.03 d/yr; *r*^2^ = 0.33, *t* = 3.30, *p =* 0.003). **(D)** Annual mean laying date advanced over time for late nesting birds (75^th^ percentile: –0.245 ± 0.09 d/yr; *r*^2^ = 0.23, *t* = 2.59, *p =* 0.016; 90^th^ percentile: –0.236 ± 0.10 d/yr; *r*^2^ = 0.19, *t* = 2.34, *p =* 0.028), but not for early nesters (25^th^ percentile: –0.033 ± 0.10 d/yr; *r*^2^ < 0.01, *t* = 0.33, *p =* 0.745; 90^th^ percentile: –0.136 ± 0.09 d/yr; *r*^2^ = 0.09, *t* = 1.49, *p =* 0.151). **(E)** Kernal density estimates (from ggplot geom_density function), which provide a smoothed version of the histogram, show longer right-hand tails in the distribution of laying date early in the study and greater constriction early and late in the distribution in later years. The five year periods contain 5 years of data each.

In contrast to the mean laying date, there has been a much larger change in the distribution of laying dates. Laying date has become 46% less variable (smaller SD) over the 26 years of study, compared with the 19% change in the mean (SD of laydate vs year: slope = –0.107 *±* 0.032, *t* = 3.33, *p =* 0.003, df = 1, 23, *r*^2^ = 0.325; Figure 2C). This has occurred because late-nesting birds are advancing laying (ie, the right tail of the distribution has shrunk) rather than changes in early-nesting. The 75^th^, 90^th^ and 95^th^ percentiles of laying date have become earlier over time (stand. slopes = –0.464 to – 0.646; Appendix S1: Table S1), while the 5^th^, 10^th^, 25^th^ and 50^th^ percentiles did not change (Figure 2D, E; Appendix S1: Table S1). This advancement occurred among older (ASY) females when age groups were analyzed separately (Appendix S1: Table S1), so it was unlikely to be related to changes in the age distribution. There were more late-nesting first-year (SY) females during the early years of the study, but <10 % of nesting females were SY after 2002. Hatching date was correlated with laying date (r^2^ = 0.98, n= 25 mean annual values) so in recent years there was also a shift toward earlier hatching among late nesting birds (Appendix S1: Table S1).

### Is the climate changing during laying or hatching?

Wind speed during the pre-laying period (1-15 May) has slowed by 47% from 1997 to 2023 (Figure 3A) and it was the only weather variable to change during the pre-laying period (Appendix S1: Table S2). Slower wind speed during pre-laying was also correlated with a narrower distribution of laying dates (smaller SD; Figure 3B) and an earlier average laying date (Figure 3C). The mean annual laying date was not correlated with any of the other weather variables (Appendix S1: Table S2).

**FIGURE 3.**
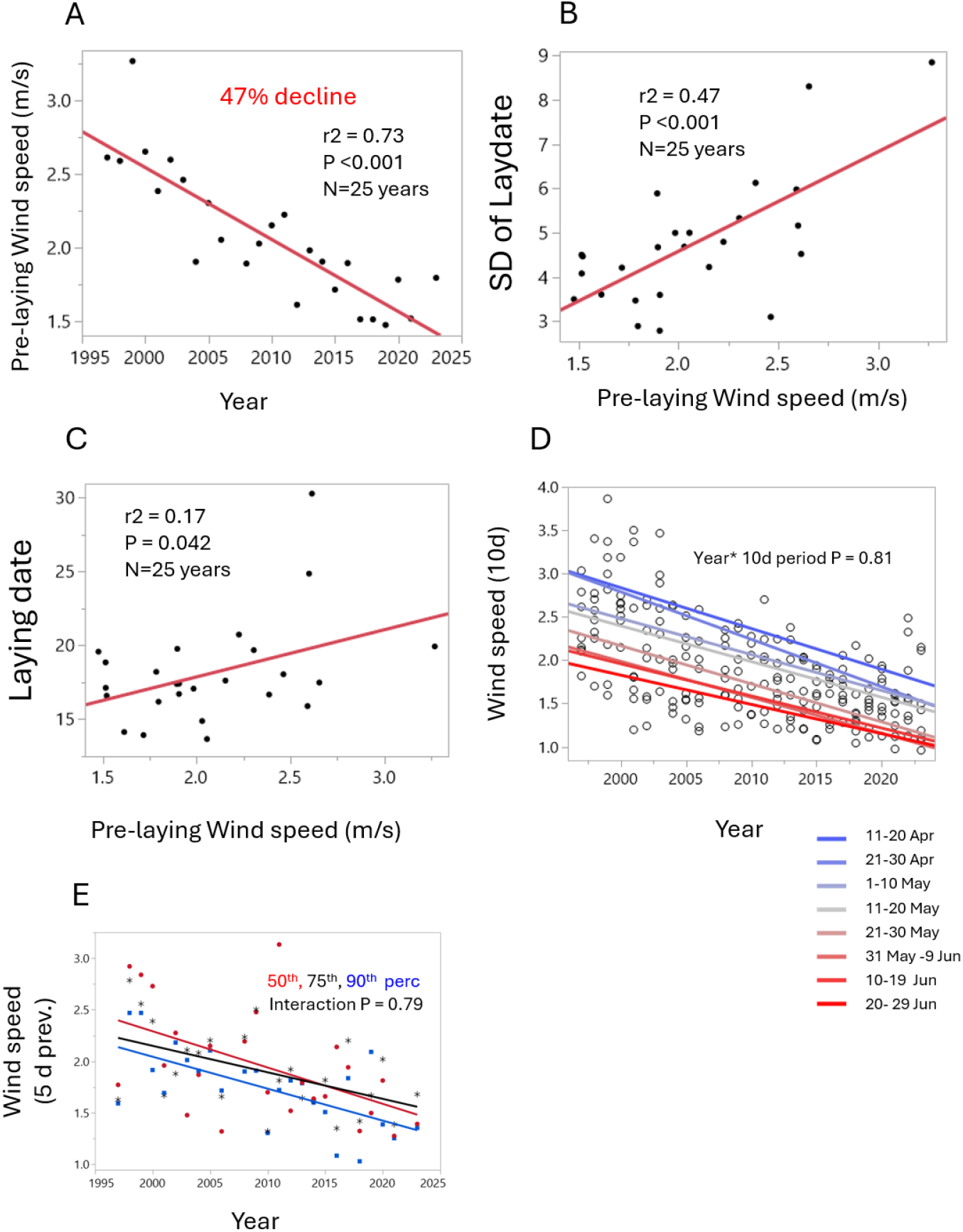
Wind speed (m/s) during the prelaying period (1-15 May) has declined 47% over 25 years in Saukville, Wisconsin, USA. (**A**; –0.049 ± 0.01 m/s/yr; *r*^2^ = 0.73, *t* = 7.9, *n* = 25 years, *p <* 0.001). Slower wind speed during pre-laying was also correlated with a smaller SD of laying dates (**B**; 2.25 ± 0.50 SD/yr; *r*^2^ = 0.47, *t* = 4.5, *n* = 25 years, *p <* 0.001) and an earlier average laying date (**C**; 3.19 ± 1.48 d/yr; *r*^2^ = 0.17, *t* = 2.2, *n* = 25 years, *p* = 0.042). Wind speed is declining at the same rate among 10 day periods from 11 Apr to 29 Jun (**D**; ANCOVA: Year *F*_1, 192_ = 29.2, *p* < 0.001; 10d period *F*_7, 192_ = 22.9, *p* < 0.001; Year * 10d period interaction *F*_7, 192_ = 0.53, *p =* 0.81). The eight 10-day periods are shown below panel **D**. The rate of wind speed slowing did not differ for females starting laying on the 50^th^, 75^th^ or 90^th^ percentiles of laying each year (**E;** ANCOVA: Year *F*_1, 69_ = 13.0, *p* < 0.001; percentile *F*_2, 69_ = 2.1, *p* = 0.136; Year * percentile interaction *F*_2, 69_ = 0.24, *p =* 0.79).

Slowing wind speed was also the only weather variable that has changed over the 26 years on the average dates for the 75^th^ or 90^th^ percentiles of laying (21 and 24 May, respectively) or the 5 or 10 d previous to those percentiles (Appendix S1: Table S2). The slowing of wind speed across years has occurred at the same rate (i.e., same slopes) throughout the breeding season when we examined 10 day periods across the breeding season from 11 Apr to 29 Jun (Figure 3D; year * 10d period interaction *p =* 0.81). Thus, the rate of wind speed slowing did not differ for females starting laying on the 50^th^, 75^th^ or 90^th^ percentiles of laying each year (Figure 3E). In other words, the decrease in wind speed (over the years) was similar for average and late nesters. Note that within a season wind speed decreased from April to June (i.e., the intercepts decreased; Figure 3D).

Climate has also changed around the time of hatch. Wind speed decreased on and around (5 and 10 days before) the long-term average date of hatch (6 June), and there was also an increase in the average air temperature and solar radiation 5 and 10 days before hatch (Appendix S1: Table S3). As with wind speed, these changes in temperature and solar radiation were similar between the median (50^th^) and late (75^th^ and 90^th^ percentile) nests (Appendix S1: Table S3).

Slowing wind speed may be a widespread pattern in southeastern Wisconsin, because wind speed has also slowed 61 km away at Milwaukee International Airport. During 1997-2021 wind speed was 9% slower during April (slope = –0.019 *±* 0.008, t = 2.46, *p =* 0.022) and 12% slower during May (slope = –0.021 *±* 0.007, t = 2.93, *p =* 0.008). There is, however, no slowing trend in most other months at the Milwaukee airport, including June (*p =* 0.56) when swallows are still breeding.

### What is changing the distribution of laying dates?

To explore the causes of long-term changes in laying dates, we switched our analyses from annual means to individual nests with Bayesian distributional mixed models. This method allowed us to examine the entire distribution (both mean and shape) and incorporate individual female effects with a random effect of individual female identity.

Prior to examining the effects of weather and food availability on laying date, we constructed a baseline model that only considered changes in the mean and shape of the laying date distribution over time. This baseline model confirmed that there has been a long-term advancement in the mean laying date (*β* syear = –1.19, 95% CrI –2.27-0.12), as well as a shift in the shape (k) of the distribution (*β* syear = 12.83, 95% CrI 2.70 –22.51; Figure 4). The Gamma shape parameter (k) is inversely related to variance (= mean^2^/k) and skew (=2/k), so larger positive values, as shown here, decrease the variance as well as the skew, resulting in a narrower distribution with a shorter right-hand tail over the 25 year study period. Importantly, Figure 4 illustrates that focusing on the mean of the distribution rather than the peak or mode can lead to opposite conclusions about the long-term trajectory of laying date. In this case, the mean laying date from the model shifted earlier by 1.9 days across the 25 years (from 137.2 to 135.3), while the mode shifted later by 2.0 days (from 131.8 to 133.8). These changes reduced the difference between the mean and mode by 71% (from 5.4 to 1.5 days).

**FIGURE 4.**
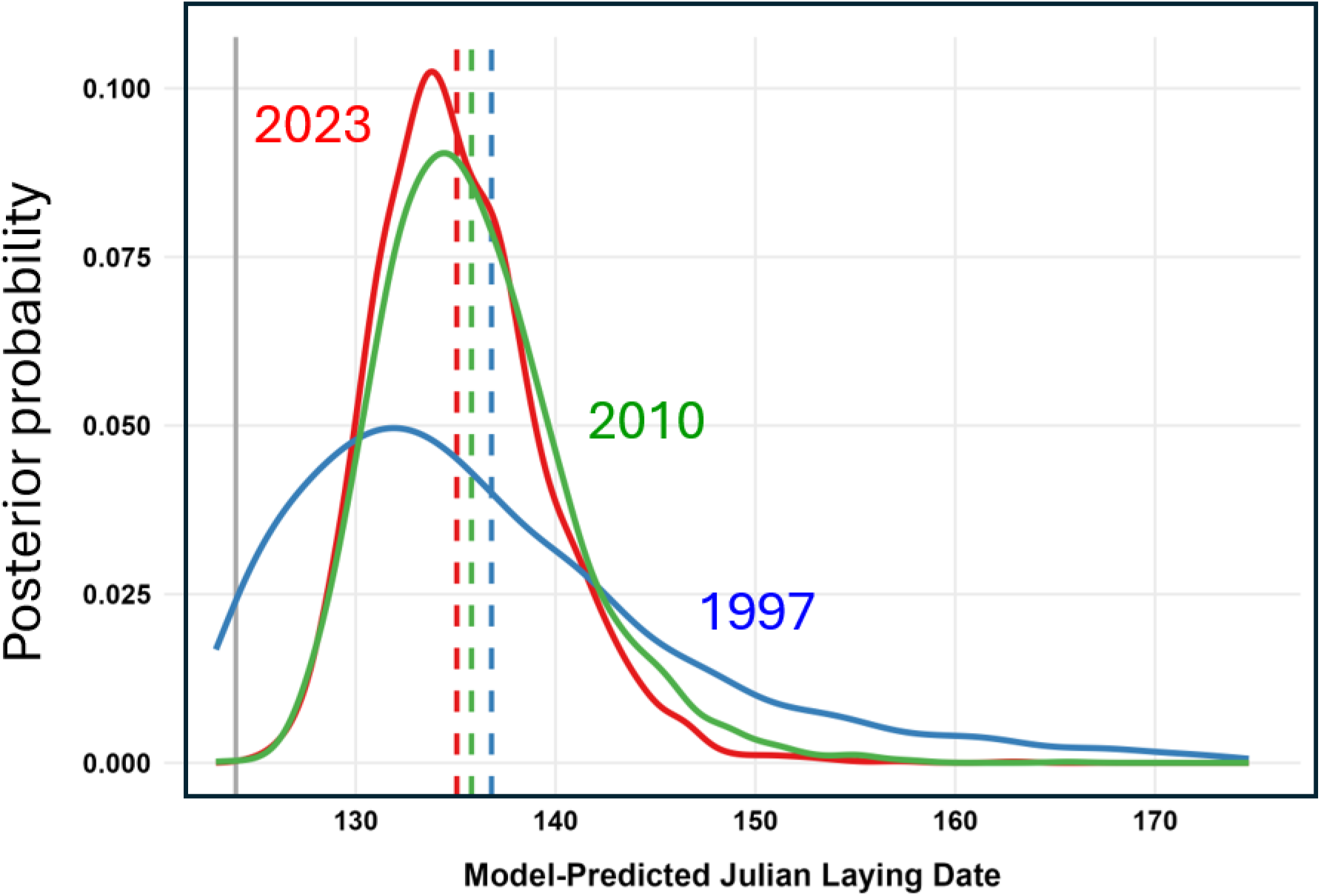
Baseline model of changes in laying date over the 25 year study period using brms to model both mean and shape of the distribution. Curves show the model predicted laying date distributions for three focal years: 1997 (blue), 2010 (green) and 2023 (red). Curves were generated using 3,000 random draws from the model’s posterior distribution. Color matched vertical dashed lines indicate the means for these three periods. The vertical gray line at Julian day 124 (4 May) indicates the earliest known laying date. Analysis of the continuous year variable indicates that there has been a long-term advancement in the mean laying date, as well as a shift in the shape (k) of the distribution (see text). Note that the mean laying date shifted earlier by 1.9 days across the 25 years (from 137.2 to 135.3), while the mode shifted later by 2.0 days (from 131.8 to 133.8).

Next, we compared this simple year-only model with several others, including models with environmental variables and those without a shape component. This comparison revealed that the highest predictive performance was achieved by a full distributional model that incorporated all of the weather and food variables and considered both the mean and shape (k) of the distribution (ELPD = 0.0; Appendix S1: Table S4). Environmental variables most strongly affected the shape, rather than the mean of the laying date distribution, so we focus on the shape first (Appendix S1: Table S5). Prelaying period rainfall had the strongest main effect on shape in the full model ( *β* = 0.73, 95% CrI 0.52 – 0.93). For illustrative purposes, Figure 5A shows that a 1.5 SD increase in rainfall above the long-term average resulted in a 198% increase in the shape parameter, which forces the distribution to become more compressed (positive values decrease variance) and, hence, laying became more synchronized in wet springs. Prelaying wind speed ( *β* = – 0.68, 95% CrI –1.03 – –0.33) and food abundance (Nematoceran biomass; ( *β* = –0.52, 95% CrI –0.78 – –0.27) were next most important, and a 1.5 SD increase in these variables led to a 64% and 54% decrease, respectively, in the shape parameter, which leads to an increase in variance (Figure 5B,C). Thus, the laying date distribution expanded when winds were calmer and there was more food (red +1.5 SD distributions in Figure 5B, C). Lastly, prelaying temperature had the weakest effect; a 1.5 SD increase in temperature resulted in a 36% decrease in the shape parameter, and thus a wider distribution of egg-laying ( *β* = – 0.30, 95% CrI –0.49 – –0.11; Figure 5D).

**FIGURE 5.**
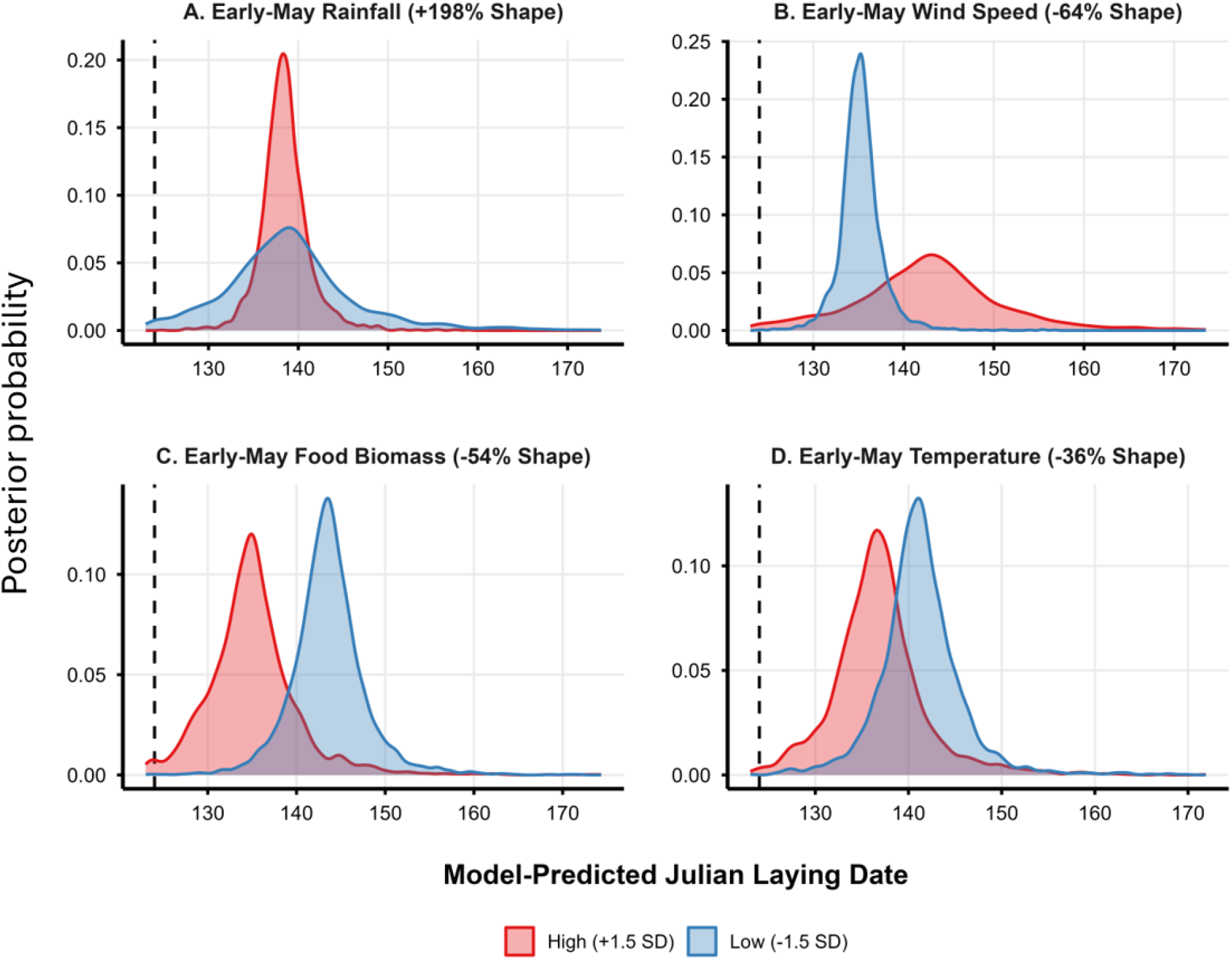
Model predicted distributions of laying date illustrating the individual effects of high (+1.5 SD) and low (–1.5 SD) rain. (**A**), wind speed (**B**), food (**C**) and temperature (**D**) during the pre-laying period. The full model results from brms are in Appendix S1: Table S5. Distributions were extracted from 3,000 draws of the full model, holding year at the study midpoint and all other environmental variables at their sample means. Effects of the interactions in the model are shown in Figure 6. Each panel contrasts high (+1.5 SD, red) and low (–1.5 SD, blue) environmental conditions for illustrative purposes. The vertical black dashed lines indicates the earliest known laying date (May 4^th^; Julian Day 124). Wet springs drive a 107% increase in the baseline shape parameter, forcing the population into a synchronized, narrow peak **(A)**. Conversely, high wind speed **(B,** –49%), food (*Nematocera*) abundance (**C**, –41%) and temperature (**D**, –26%) decrease the baseline shape parameter, allowing the population distribution to stretch and expand into an elongated right-hand trailing tail.

The strongest effects on the shape of the laying date distribution came from two interactions; one between wind and food, and one between wind and year (Appendix S1: Table S5). The negative interaction between wind and food ( *β* = –0.71, 95% CrI –0.98 – – 0.45) indicated that laying date compresses more under both calm winds and high food levels. This is illustrated in Figure 6 which contrasts the compressed laying date distribution under these good conditions (Figure 6A) with the broader distribution when higher winds decrease food availability and the distribution spreads out (Figure 6B). The model also revealed a strong positive interaction between study year and prelaying wind speed ( *β* = 0.29, 95% CrI 0.09 – 0.50), which explained over half of the total variance in the shape of the laying date distribution (Bayesian R^2^ = 0.501; 95% CrI: 0.500, 0.503). In this case, the 25-year slowing of wind speed has led to a more synchronized bottleneck with more birds laying early (Figure 6C,D).

**FIGURE 6.**
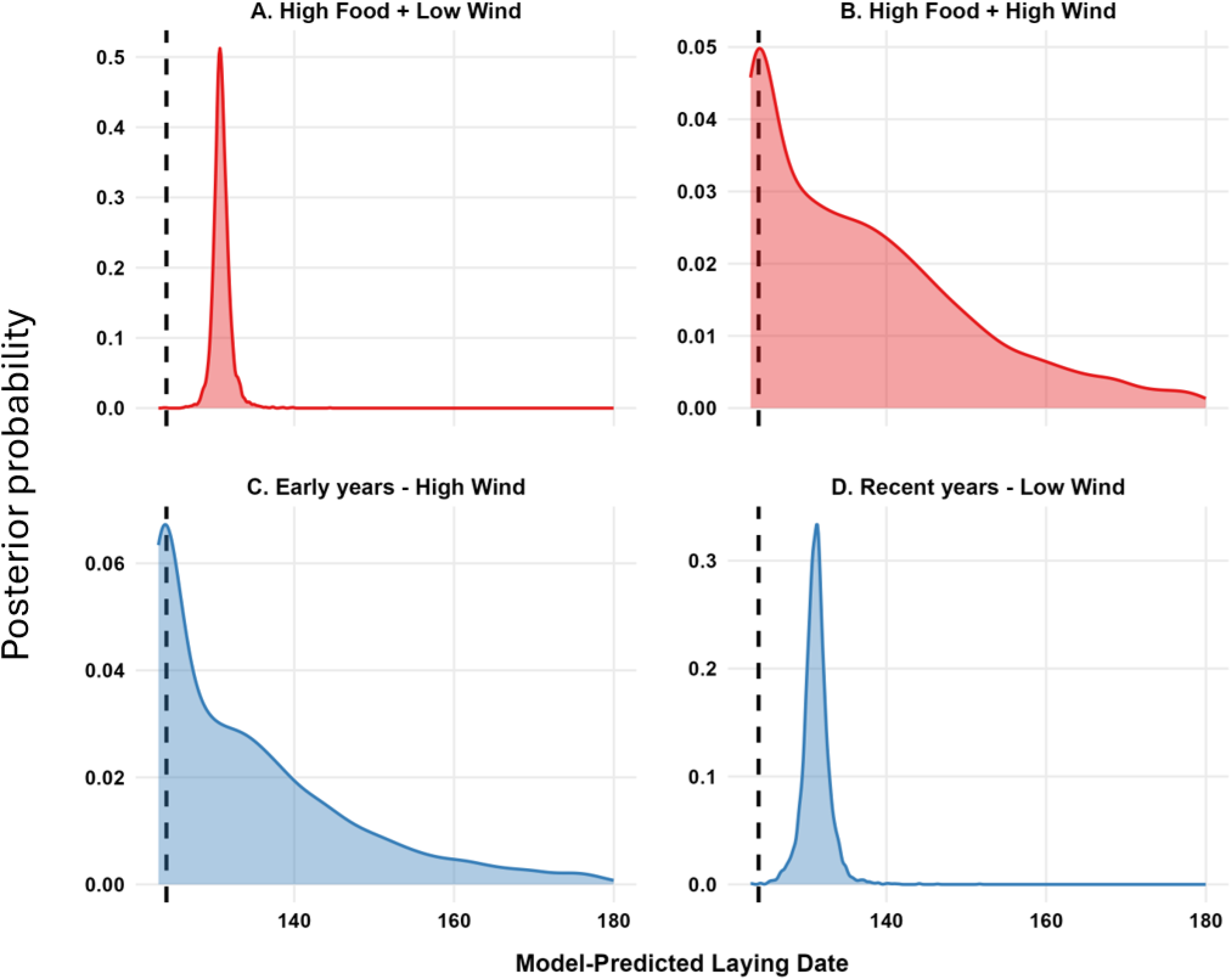
Model predicted distributions of laying date illustrating the interactions between wind speed and food (top row) and wind speed and year (bottom row) from the full model results in Appendix S1: Table S5. To illustrate the interactions, each scenario compares laying distributions under high food conditions (+1.5 SD) with varying wind speed or year (*±*1.5 SD from the mean), while holding other variables at their long-term average. Thus, the top row shows highly synchronized laying dates under calm conditions (**A,** –1.5 SD wind), but delayed breeding and a longer right-hand tail (**B**) when it is windy (+1.5 SD) and foraging is presumably more difficult. In the top panels year is held at the midpoint of the study (2010, scaled year = 0). The bottom row illustrates (**C**) the broader distribution of laying in the early years of the study (–1.5 SD for scaled year, approximately 1998) when there were higher wind speeds (+1.5 SD) versus (**D**) the highly synchronized distribution in more recent years (+1.5 SD for scaled year, approximately 2022) with calmer winds (–1.5 SD).

Turning to the mean section of the full model, the mean laying date was mainly influenced by food and wind; rain in the prelaying period had non-significant effects ( *β* = – 0.02, 95% CrI –0.05 – 0.01; Appendix S1: Table S5). One of the strongest effects on mean laying date was prelaying wind speed which pushed laying date later when winds were stronger ( *β* = 0.18, 95% CrI 0.12– 0.24); a 1 SD increase in wind speed resulted in a 20% delay in the average laying date. There was also a positive interaction between prelaying wind speed and food (*Nematocera*) biomass ( *β* = 0.08, 95% CrI 0.06 – 0.11), which indicates that high prelaying wind speeds opposed the advancement of laying date when there was more food ( *β* for the main effect of food = –0.17, 95% CrI –0.21 – –0.14). Mean laying date has actually been delayed slightly in more recent years (year *β* = 0.07, 95% CI 0.003 – 0.13) after controlling for these environmental variables and interactions, because there has been a change in the shape of the distribution. More birds are piling up at the beginning of the season producing a slightly later mean (e.g., compare Figure 6C, D). Overall, the model explained 35% of the variance in mean laying date (Bayesian R^2^_mu_ = 0.353, 95% CrI: 0.327-0.376).

### Are there constraints on breeding earlier?

The lack of change over the years in early (e.g., 5th, 10^th^, 25^th^ percentiles) and mean laying dates suggests that there might be a constraint on birds laying too early in the season. The average date of the first laid egg was 8 May and the earliest egg of any nest was laid on 4 May. A changepoint analysis indicated that both mean and minimum temperature shifted on 29 –30 April (only 1 change detected for each; Appendix S1: Table S6). Comparing the final five days of April to the first five days of May, mean temperature rose by 2.8°C and minimum temperature rose by 2.5°C. These were the highest temperature increases between any consecutive five-day blocks across the entire breeding season (average = 0.82°C–0.84°C change; Appendix S1: Figure S1).

Similarly, Nematoceran biomass showed two changes, first on 3 May and then on 20 May. The increase on 3 May averaged 0.90 mg/day (back-transformed from Ln) and was the second largest transition between 5 day periods during the breeding season (average = 0.07 mg/day increase; Appendix S1: Figure S1). The largest increase (1.3 mg/day) occurred during a second changepoint for Nematoceran biomass on 20 May, which was near the average laying date (18 May; Appendix S1: Figure S1). Combined, these temperature and food changes indicate a large shift in environmental conditions at the transition from late April to early May.

### The threshold for early breeding has been stable

The analysis above focused on shifts in the environmental variables, but we also wanted to determine if the start of the breeding window (5^th^ percentile) has remained relatively fixed or is shifting over time. Quantile regression analysis indicated that the 5^th^ percentile of laying advanced by 1.8 days over 21 years (2000-2021) in a full model (*β* year_scaled = −1.28, SE=0.56, *p =*0.023; Appendix S1: Figure S2, Table S7) that contained our weather and food variables and interactions, similar to the full distributional model (Appendix S1: Table S7. However, incorporating the interactions in the full model provided a small 2.7% gain in variance explained (Δ Pseudo-*R*^2^ = 0.027; Appendix S1: Table S7), and without them the year trend was not significant (*β* year_scaled =−0.68, *p =*0.378; Appendix S1: Table S7). Thus, while the overall distribution of laying dates has been compressing inward over time, the lower (5^th^ percentile) boundary has remained relatively stable.

### Is the food supply changing?

#### Insect Biomass

Wind speed has slowed (Figure 3), and nematoceran biomass was negatively related to wind speed (Figure 7A), so we might expect an increase in food abundance over time. Indeed, there was an increase in Nematoceran biomass from 2000 to 2021, but it occurred primarily before 2008. There has been relatively little change in food abundance since 2008 (Figure 7B; year group *p <* 0.001). The long-term increase in nematoceran biomass has also occurred primarily later in the breeding season (Figure 7C), when wind speed is slower (Figure 3D). Over the long-term, biomass has remained relatively stable early in the season, during the pre-laying (1-15 May) and laying periods (25^th^ to 75^th^ percentiles of laydates; *p =* 0.57), but biomass has increased later in the season during the incubation (*p =* 0.02) and nestling periods (*p =* 0.04; Figure 7D).

**FIGURE 7.**
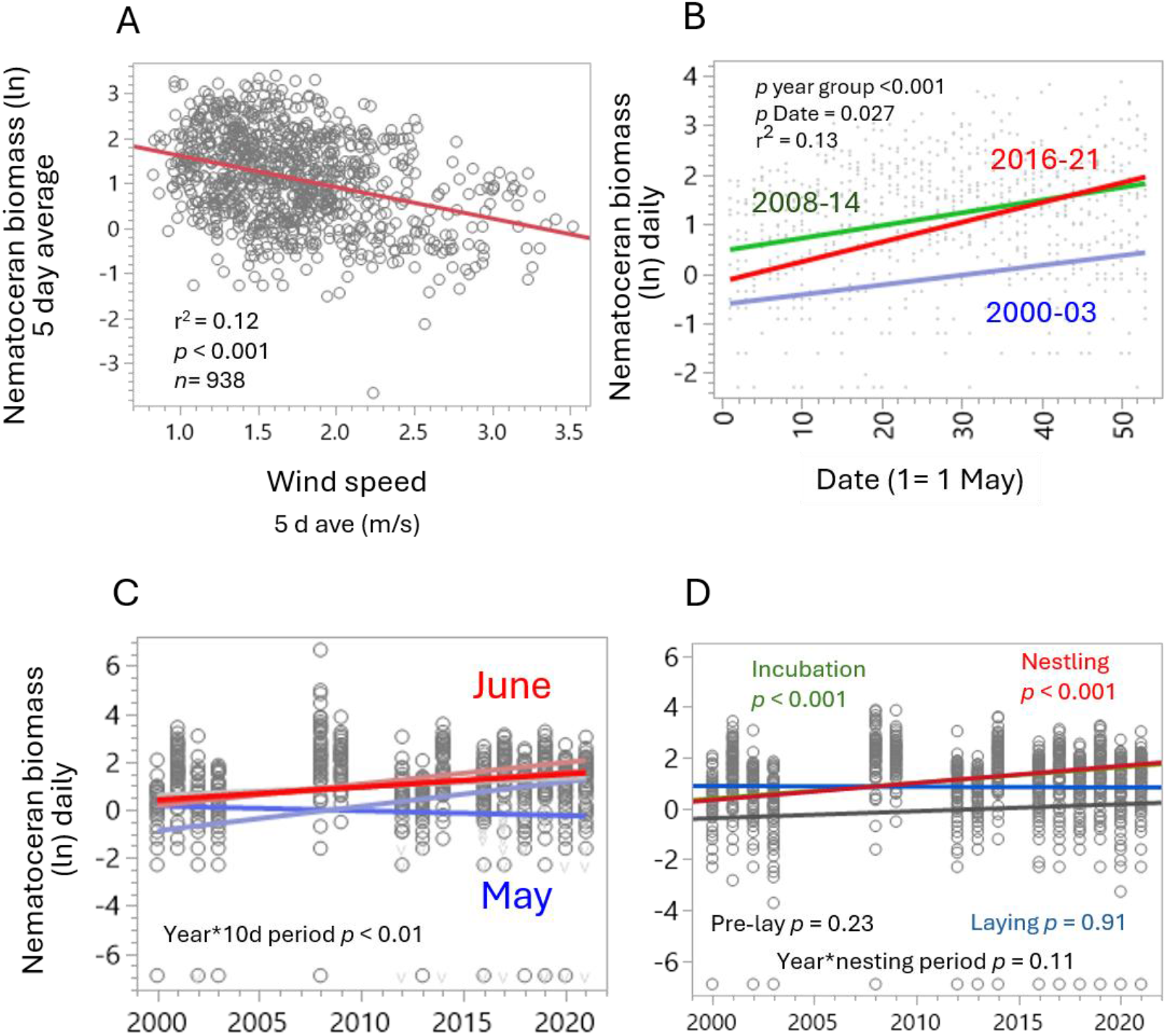
Food abundance (Nematoceran biomass, mg, ln-transformed + 0.001) in relation to wind speed. (**A**), date in the season (**B**) and year for 10 day periods starting in May (**C**) and for stages of nesting (**D**). **A**. Nematoceran biomass declined with stronger wind speeds (*r*^2^ = 0.12, *p* <0.001, *n* = 939; five day averages were used to smooth out extreme values). **B**. Daily nematoceran biomass increased across the season (May to June; F_1,972_ = 4.57, *p =* 0.033) and was higher in middle (2008-14) and late (2016-21) than in the early years (2000-03) of study (F_2,972_ = 22.9, *p <* 0.001); the interaction between day and year group was not significant (F_2,972_ = 1.82, *p =* 0.16). Overall *R*^2^ = 0.13. **C**. Daily nematoceran biomass increased across the season using 10 day periods (blue to red shading indicates May to June; F_5,822_ = 9.48, *p <* 0.001) but the difference was more extreme in recent years (year* 10d period interaction; F_5,822_ = 3.13, *p <* 0.01). Overall *R*^2^ = 0.11. **D**. Annual nematoceran biomass also increased across years F_1, 769_ = 12.6, *p <* 0.001) in an ANCOVA with four nesting periods (year*nesting period, F_3, 769_ = 1.98, *p =* 0.11). Nematoceran biomass increased during the incubation (green line hidden under nestling line; F_1, 231_ = 3.62, P<0.001) and nestling (red line; F_1, 258_ = 27.2, P<0.001) periods, but not during the earlier pre-laying (black line; F_1, 215_ = 1.41, P= 0.24), or laying (blue line; F_1, 65_ = 0.01, P= 0.91) periods. The interaction between year and these nesting periods was not significant (F_3, 769_ = 1.98, P= 0.11).

#### Timing of peak food abundance

The peak date of nematoceran biomass (based on a 5 day running average) has not changed over the 22 year span of insect sampling (slope = –0.579 *±* 0.516, *n* = 15 years, *t* = 1.12, *p* = 0.283, *r*^2^ = 0.09). The average date for the peak of nematoceran biomass was 4 June (3.66 se, 95% CI: 27 May – 11 June, *n* =15 years). Averaged across all the years, nematocerans showed a gradual increase in biomass through May and then a slight decline toward the peak of hatching (6 June, Figure 8A). However, there was substantial yearly variation as shown in Figure 8B, which averages Nematoceran biomass during three time periods of the study (early, middle and late years), and in Figure 8C for each year within those time periods. There was no relationship between the average dates of laying (slope = –0.048 *±* 0.045, t = 1.06, *p =* 0.307, *n* = 15 years) or hatch (slope = –0.049 *±* 0.051, *t* = 0.97, *p =* 0.351, *n* = 15 years) and peak date of nematoceran biomass.

**FIGURE 8.**
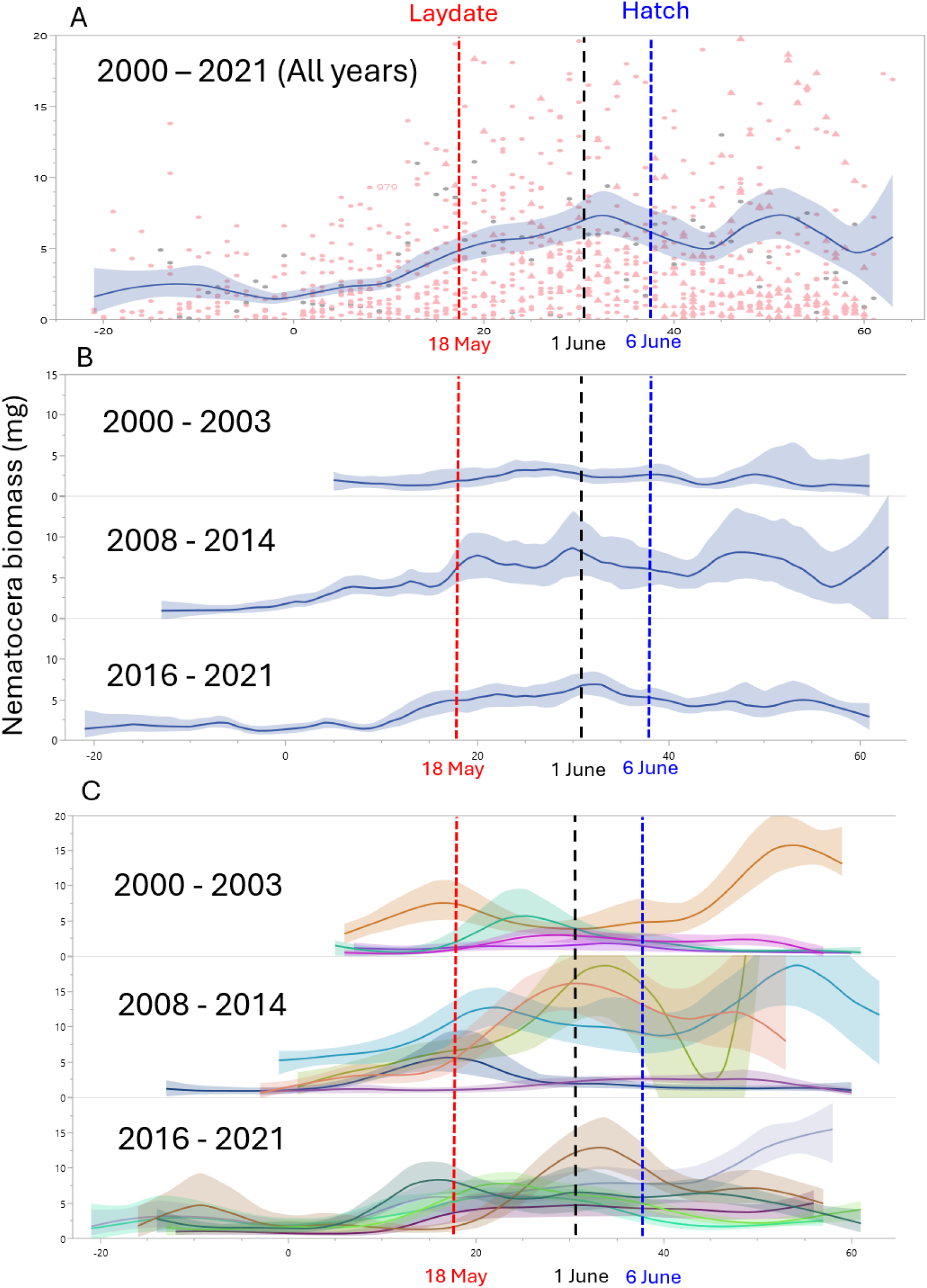
Nematoceran biomass (mg/day) in relation to date (1= 1 May) averaged across all years. (**A**), three sequential time periods with (**B**), or for individual years within the three time periods (**C**). Vertical dashed lines indicate average laying date (18 May), 1 June and average hatching date (6 June). Lines are splines with confidence bands. There were 15 years of insect sampling using a suction trap across a 22 year span (2000-2021).

### Is there selection against mismatched laying?

The peak of Nematoceran biomass averaged 2.2 days (*±* 3.8 se, *n* = 15 years) before the average hatch date, and the difference between these dates (the mismatch) has not changed in the 22 years spanning 2000 and 2021 (b = –0.506 *±* 0.548 se d/yr, *r*^2^ =0.062, t= 0.92, *p =* 0.373, *n* = 15 years with suction trap data).

Although the date of peak biomass has not shifted consistently over the years, it has varied considerably (interquartile range = 16 days before to 15 days after hatch; Figure 9). Indeed, there was no clear time in the season for the peak in biomass. In six of the 15 years of insect sampling, the peak of nematoceran biomass was closer to the average laying date for that year (2.6 days after laying *±* 1.557 se) than to the average hatch date (16.5 days before hatch *±* 1.506 se). However, in another four years, the nematoceran peak was closer to the mean hatch date for that year (4.2 days before hatch *±* 1.194 se; 1.2 – 6.9 d before) than to the laying date (15.2 *±* 1.305 d). In the remaining five years the peak of nematocerans was closer to fledging (range: 14 to 21 days after hatch) and the end of our insect sampling period (23 June; Figure 9).

**FIGURE 9.**
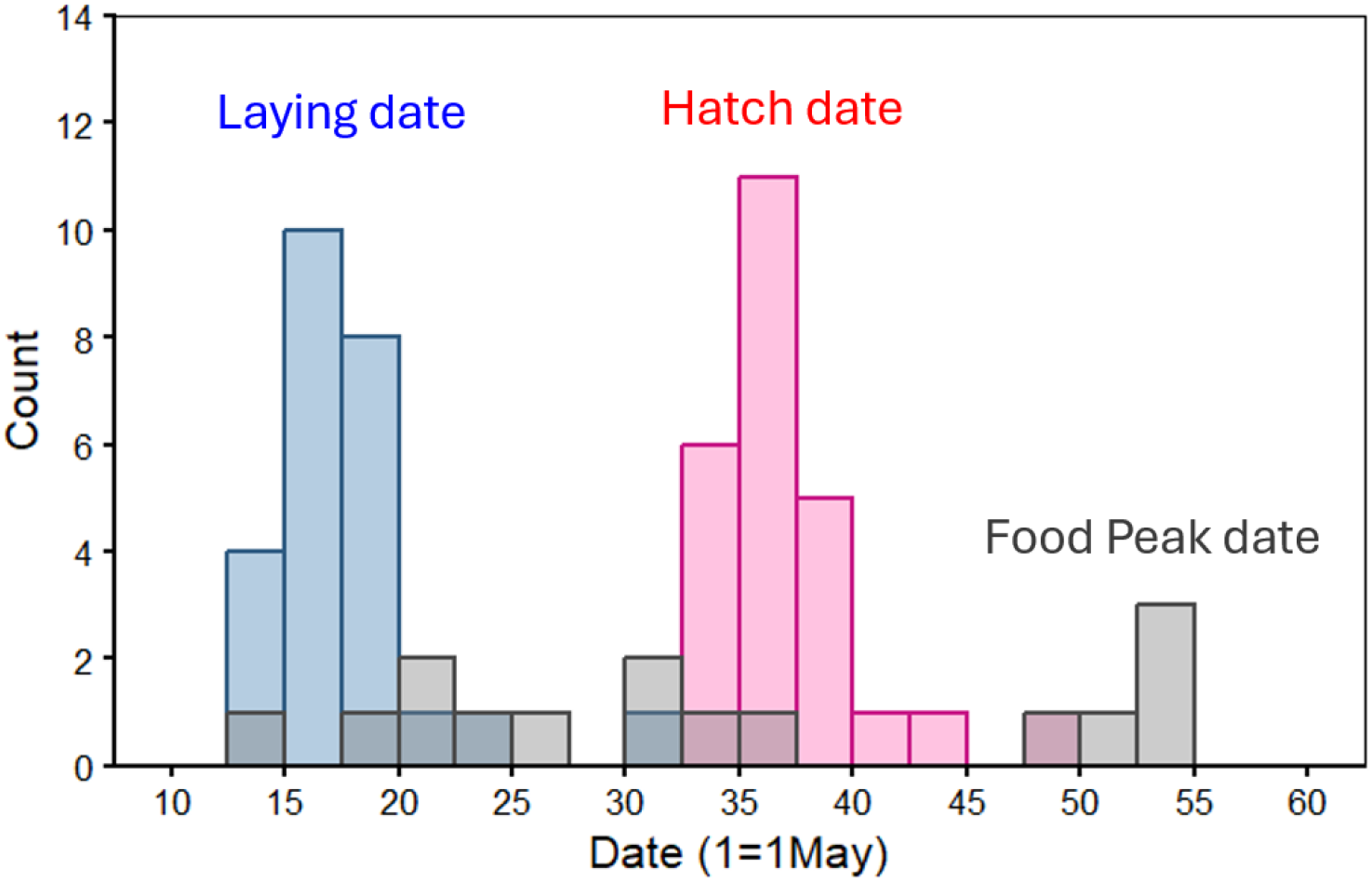
Distribution of annual means for laying and hatch dates and peak date of food (Nematocera) biomass. Peak food date was calculated separately for each year from a running 5 day average.

Despite this variability, the annual date of peak biomass was significantly related to all of the weather variables in a sliding window analysis (i.e., climwin; Appendix S1: Table S8), and the peak biomass date could be predicted accurately (R^2^ = 0.84) with the two strongest predictors, maximum temperature (maxT) during 18-30 Apr (4.12 *±* 1.01, t = 4.1, *p <* 0.002) and mean rainfall during 1-26 April (5.38 *±* 1.29, t= 4.2, P<0.002; df = 2, 12, multiple regression; Appendix S1: Table S8). Thus, the date of peak nematoceran biomass is potentially predictable based on weather conditions in April, at least a week before the first eggs are laid (10 May is the average date for the 5^th^ percentile of laying dates; Figure 1). Note that most birds have settled at a nestbox and started nest building by mid-to late-April and are exposed to these weather conditions.

#### Selection on mismatching

Mismatching with the peak of food abundance produced no overall directional (linear) selection (Figure 10A, Appendix S1: Table S9; beta = 0.01, 95% CrI –0.07, 0.09; probability of direction = 59.9%); however, there was strong stabilizing selection, particularly at the beginning of the study (Figure 10B, Appendix S1: Table S9; beta = –0.08, 95% CrI –0.17 –0.00; probability of direction = 97.9%). The stabilizing selection was equivalent to a Lande-Arnold stabilizing gradient (gamma) of –0.16. However, stabilizing selection has declined over the course of the study (Figure 10 B, Appendix S1: Table S9). The interaction between the quadratic mismatch term and decade (year/10) was positive (beta = 0.06, 95% CrI 0.00 –0.13; gamma = +0.12 per decade), and it produced a high Bayesian probability of direction (pd = 97.6%) for the flattening of the curve. Female age class also had a significant effect with younger (SY; beta = –0.11, 95% CrI –0.23 – 0.00; pd = 97.2%) and unknown age (beta = – 0.10, 95% CrI –0.18 – –0.02; pd = 99.6%) individuals experiencing lower relative success than presumably older (ASY) females. This suggests that the temporal shifts in selection were not due to demographic changes in age structure.

**FIGURE 10.**
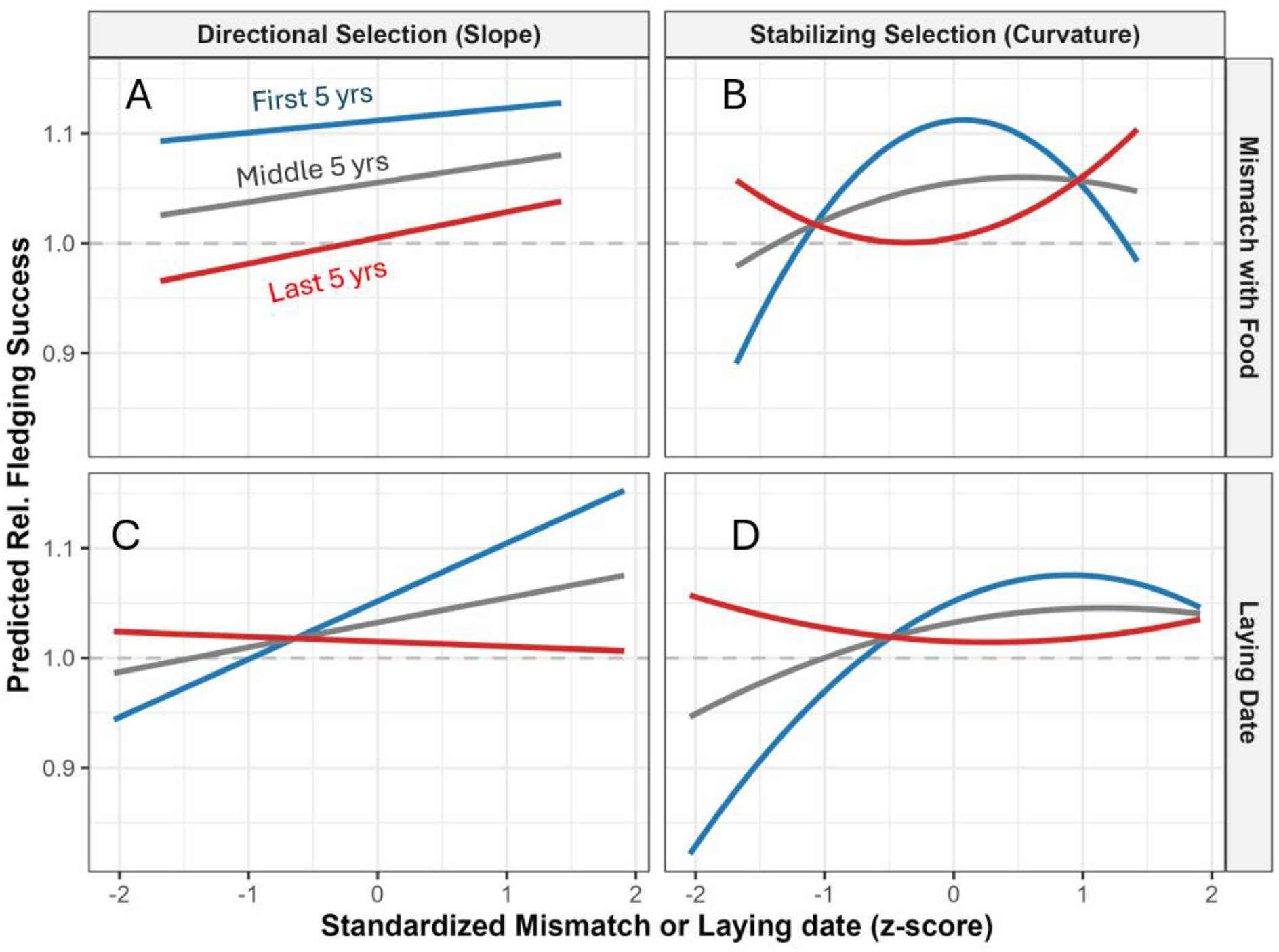
Long-term directional (left column) and stabilizing (right column) selection on mismatching (top row; N= 932 nests) and laying date (bottom row; n = 1,351nests) over a 22-year study period. Average relative fledging success (w, scaled to 1.0) is indicated by the horizontal dashed light-gray line and is plotted against standardized phenotypic values (z-score; mean = 0, SD = 1) for mismatch or laying date. Panels show three time periods (“First 5 Years,” solid blue line; “Middle 5 Years,” solid gray line; “Last 5 Years,” solid red line). The top row shows weak (probability of direction = 59.9%) linear selection (**A**) and strong (pd = 97.9%) stabilizing selection on mismatching (**B**). However, stabilizing selection is weakening over time (pd = 97.6%); so the convex red line in B (last 5 years) was not significant (pd = 84.3%). The bottom row shows both linear (probability of direction = 95.6%; **C)** and stabilizing (pd = 96.2%; **D**) selection on laying date. However, these trends also weakened over time (linear pd = 88.9%; stabilizing: pd = 91.1%). Predictions hold female age constant at the reference class (After Second Year) and control for annual variation using a Bayesian multilevel model with random intercepts and slopes.

The weakening stabilizing selection can be seen by comparing three time periods of the study (first 5 years, middle and last 5 years; Figure 10B). Note that although the shape of the stabilizing selection curve seemed to change to an upward convex curve during the last 5 years, the quadratic estimate (beta = 0.032, 95% CrI –0.0288 – 0.0989) overlapped zero and the probability of directional certainty was weak (pd = 84.3%), indicating that by the end of the study period there was no significant selection occurring due to mismatching.

#### Selection on laying date

In contrast to selection from mismatching, selection on laying date showed both directional and stabilizing selection overall, but it also declined over time (Figure 10C, D). This was supported by strong Bayesian probability of direction (pd) values (>0.95) for the entire study period, despite 95% credible intervals (CrI) that marginally overlapped zero (Appendix S1: Table S10). Overall, there was moderate stabilizing selection (beta = –0.03, 95% CrI –0.07 – 0.00; Lande-Arnold gamma = –0.06; probability of direction = 96.2%) and positive directional selection favoring delayed breeding (beta = 0.06, 95% CrI –0.01-0.13; probability of direction = 95.6%). Often there is directional selection for earlier laying, but these results indicate that there was greater reproductive success when the entire stabilizing function was shifted later in the season (due to the directional selection). However, this pattern weakened over the course of the study. The probability of direction (pd) for a long-term decline in linear selection was 88.9% (Figure 10C; beta = –0.03, 95% CrI –0.09 –0.02; Table S9) and 91.1% for a decline in stabilizing selection (Figure 10D; beta = 0.02, 95% CrI –0.01 – 0.05). These trends can be seen by comparing the estimates across the three 5 year time periods (Figure 10C, D). As with the mismatching analysis, female age class had a significant effect with younger (SY; beta = –0.11, 95% CrI –0.24 – 0.02; probability of direction = 95.1%) and unknown (beta = –0.05, 95% CrI –0.10 – 0.00; probability of direction = 96.1%) age individuals experiencing lower relative success than presumably older (ASY) females.

### Does weakening selection and wind speed affect fledging success?

Next we wanted to explore the potential effects of weakening selection and wind speed on trends in other aspects of reproduction, such as fledging success. First, we examined clutch size, which has been increasing over time (*p =* 0.009, r2 = 0.26, *n* = 25 years, Figure 11A) and is negatively related to wind speed prior to laying (1-15 May, *p =* 0.002, r2 = 0.34, *n* = 25 years; Figure 11B). Thus, the decrease in wind speed over time was also associated with an increase in clutch size. Fledging success (Figure 11C) and the mean weight of nestlings (measured on day 12; *p <* 0.001, Figure 11D) have also increased over time. The increase in nestling weight was also related to decreasing wind speed (*p =* 0.037, Figure 11E) which, in turn, was related to greater food abundance during the main period of nestling growth (days 1-12; *p =* 0.021, Figure 11F).

**FIGURE 11.**
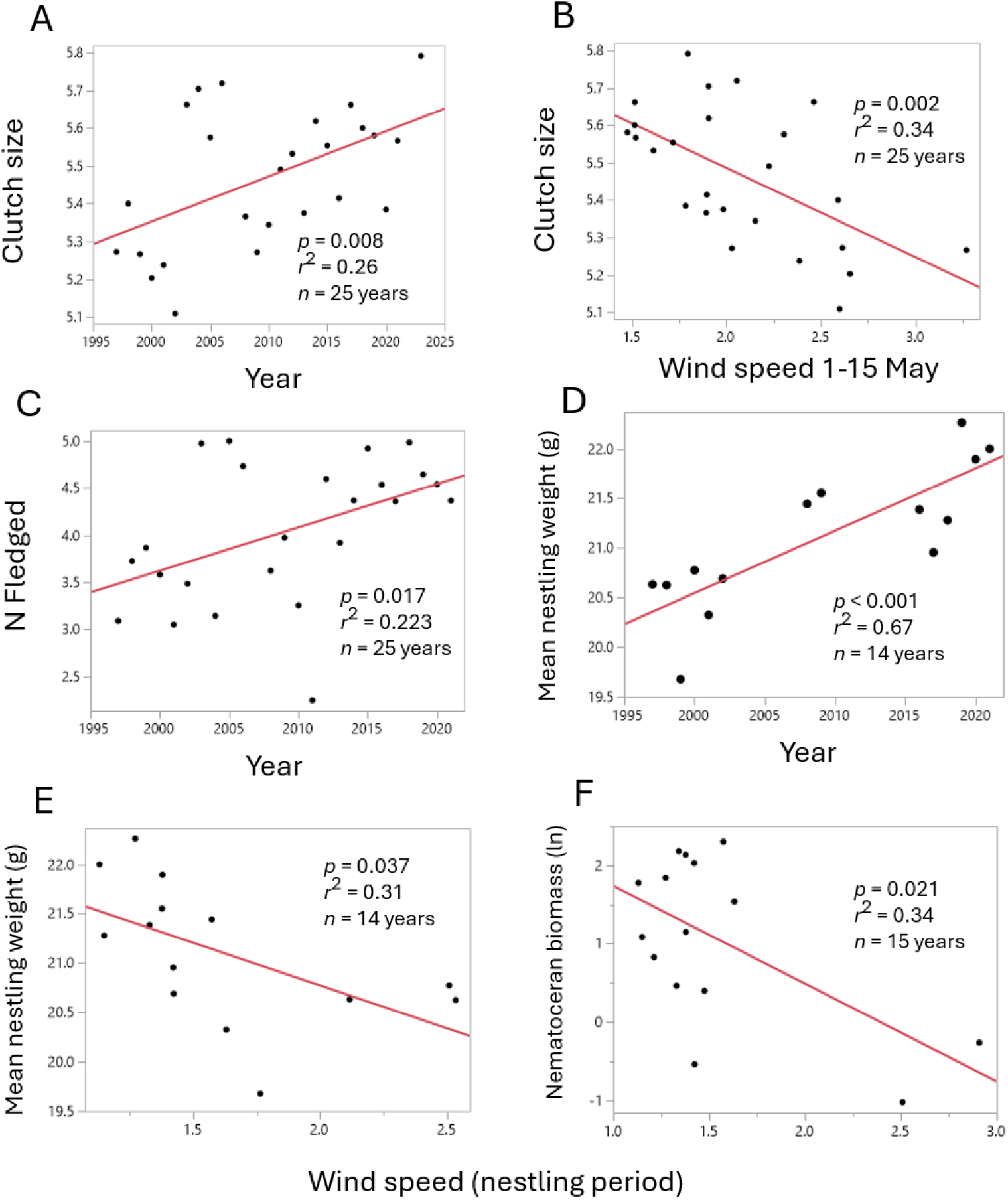
Clutch size increased over time. (**A**) and it was negatively related to wind speed during the pre-laying period, 1-15 May (**B**). Fledging success (**C**) and the mean weight of nestlings (measured on day 12; **D**) also increased over time. The increase in nestling weight was also related to decreasing wind speed (**E**) which, in turn, was related to greater food abundance during the main period of nestling growth (days 1-12; **F**).

### How do food, temperature and wind affect reproductive success?

Food abundance, temperature and wind can interact and affect different stages of reproduction after laying, from clutch size to the number of young that fledge. To explore these relationships, we constructed a structural equation model (SEM; see Methods and Appendix S1) to test the relative importance of each environmental variable and the timing of its potential effect. Test results are shown from JMP, but they were nearly identical to those from the R packages lavaan (Rosseel, 2012) and piecewiseSEM (Lefcheck, 2016; Appendix S1)

Surprisingly, the initial (full) SEM indicated that there was no significant effect of food abundance at any stage after the pre-laying period (dashed lines in Figure 12A, Appendix S1: Table S11, 12). Only food biomass pre-laying had an effect on the reproductive variables. To simplify the model (Figure 12A), we removed the post-laying food variables, as well as two relatively weak paths (B’ <0.1) between: 1) wind speed and the number of nestlings on day 12 (b’ = –0.069), and 2) temperature during the nestling period and the number of fledglings (b’= –0.049).

**FIGURE 12.**
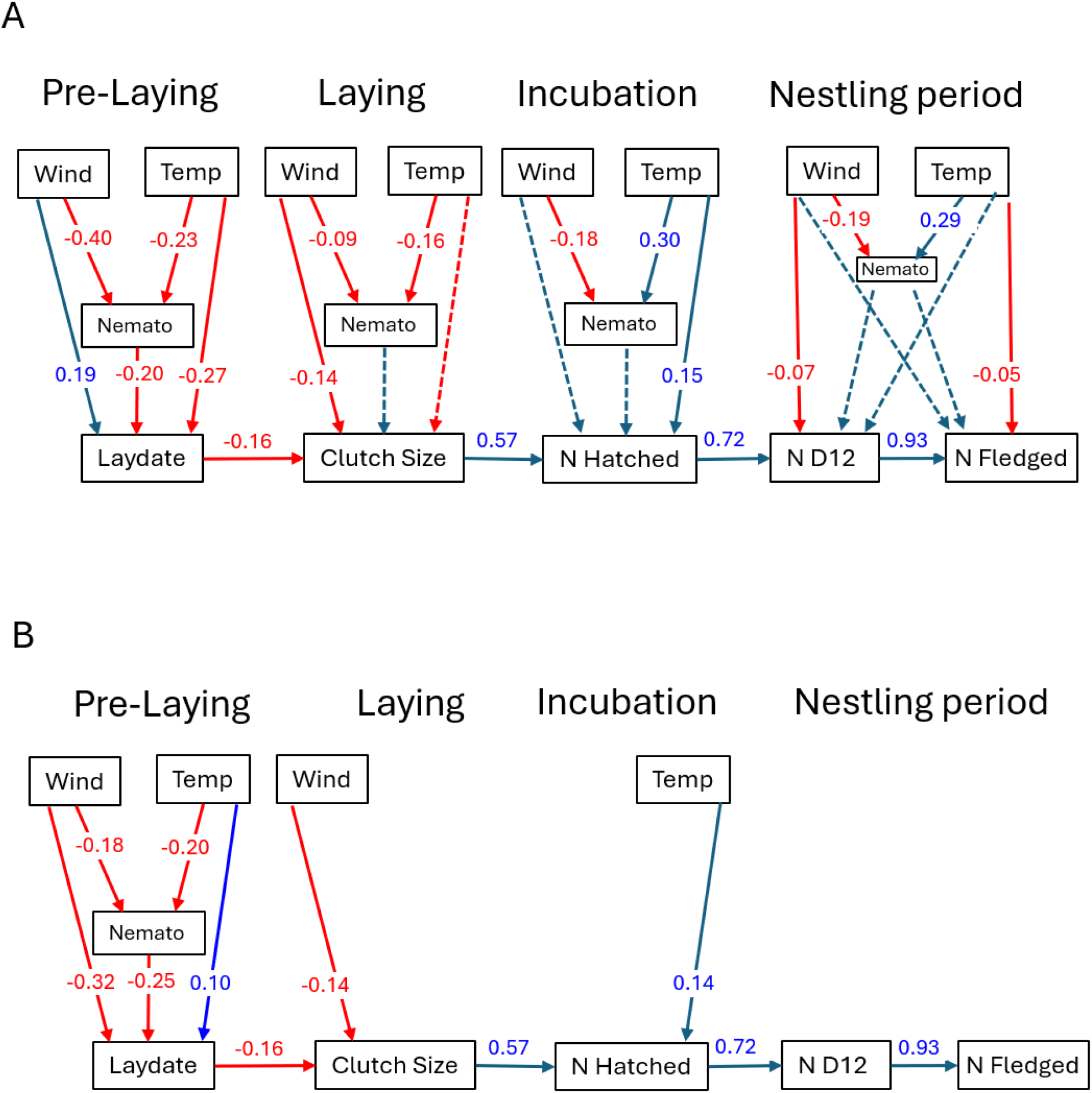
Structural equation models (SEM) of reproductive success in relation to food abundance (Nematocera biomass), air temperature and wind speed. The initial full model (**A**) and the final reduced model (**B**) with only significant paths with *b*’ >0.1 were included. Numbers on paths are standardized path coefficients. Red arrows and numbers indicate negative relationships, and blue arrows and numbers indicate positive relationships. Solid lines were significant (*p* <0.05), dashed lines were not significant and were removed for panel **B**. See supplemental materials for more details and Appendix S1: Table S13 for a comparison of results from different methods.

The final (simplified) model (Figure 12B) indicated three main results. First, only food abundance during the pre-laying period had effects on laying date (*p <* 0.001) and later stages of reproduction, including clutch size (P= 0.002) and number of young fledged (*p =* 0.002). Surprisingly, food abundance during the nestling period, for example, did not have a significant effect on the number of young surviving to day 12 or the number that fledged, once the pre-laying effects of food on clutch size were taken into consideration.

Second, warmer temperature during incubation was positively related to the number of eggs that hatched. Hatching failure accounts for one of the largest losses of reproductive success. Half of nests (50%, 273/548) had some loss of eggs or young, and hatching failure was the largest cause (21 ± 1.6 SE % of eggs did not hatch), followed by loss during the early nestling period (17 ± 1.7 SE% of young were lost during days 1-12) and late nestling period (6.5 ± 1.0% of young were lost after day 12). Temperature during incubation was also the only significant predictor of nesting success (binomial) among 11 weather and food variables during the incubation or nestling periods (False discovery rate *p <* 0.001; Appendix S1: Table S13).

Lastly, we found that wind speed had multiple effects on laying date and clutch size, including direct and indirect (through food) effects on laying date (*p <* 0.001) and direct effects on clutch size (b’ = –0.136). Higher wind speed decreased nematoceran biomass which delayed laying (the indirect effect) as well as directly delaying laying (direct effect of b’ = 0.103). There was only a direct negative effect of wind speed on clutch size (b’ = 0.136), so increasing wind was associated with smaller clutches, but this did not appear to occur through an indirect pathway via food, which was not significant (Figure 12A, b’= –0.002, *p =* 0.584), and, hence, it was removed in the final model (Figure 12B).

### Differing effects of food abundance on nesting success in NY and WI

These results from the SEM appear to differ from those found in Ithaca NY, where Nematoceran biomass during nestling days 7-12 was positively related to several measures of fledging success (Twinning et al., 2018). To examine this apparent difference, we performed similar analyses using data from our Wisconsin population. As in the NY study, we found that Nematoceran biomass during nestling days 7-12 was positively related to the number of fledglings (b’ = 0.10, *p =* 0.03) in a model that included hatch date and the number of cold days during nestling days 7-12. However, adding clutch size or the number of eggs that hatched (brood size) to the model led to a non-significant effect of Nematoceran biomass on fledging success. Thus, the results became non-significant after accounting for the baseline number of young (eggs or chicks that hatch) as in our path analysis. Fledging success (N fledged/N hatched from successful nests), allows one to focus specifically on changes in reproductive success during the nestling period, and it was also not related to Nematoceran biomass during 7-12 d (or any other 6 day period) in our WI population. In contrast, fledging success in the NY population was positively related to Nematoceran biomass during 7-12 d of age. Similarly, nesting success (whether or not a nest produced any fledglings, coded as 0 or 1), was not related to nematoceran biomass in WI, but it was positively related to biomass in NY. Thus, when we use similar analyses, there appear to be differences between WI and NY in the effect of Nematoceran biomass on fledging and nesting success.

## DISCUSSION

Many ecological studies of climate change focus on the effects of increasing air temperature on the average timing of reproduction and its consequences for reproductive success. At our study area in southeastern Wisconsin, USA, there was no significant change in mean air temperature during 1997-2023, nor was there any change in the mean laying date of Tree Swallows. Thus, in our case, climate and the timing of breeding did not appear to be changing, if one focuses only on temperature and the annual means of laying date. However, there has been a dramatic (47%) decrease in wind speed, and birds have responded with a sharp reduction in the variance of laying dates, rather than a change in the annual mean date. Laying date has become less variable because: 1) early nesting birds cannot advance further due to temperature and food constraints early in May, and 2) late nesting birds are advancing due to the increasing food availability caused by slowing wind speed. Slower wind speed was associated with greater availability of food (Figure 6A), primarily later in the season (Figure 6C, D), as well as an increase in clutch size, heavier nestlings and more fledglings (Figure 11). Based on a structural equation model, we found that food and weather early in the season (pre-laying and incubation) had the strongest effects on fledging success, rather than environmental conditions later during the nestling period. This result was surprising because many previous studies of birds have found that food and temperature during the nestling period have the greatest influence on reproductive success. This is particularly important to the mismatch hypothesis because it focuses on the energetic needs of developing offspring (i.e., in contrast to producing the eggs). However, selection from mismatched timing of breeding was only important early in the study period and it rapidly declined in importance with the dramatic decline in wind speed. Overall, these results illustrate that understanding how climate change is affecting reproductive success requires detailed information about the influence of food and weather on the full distribution of laying dates and their subsequent effects through to fledging. Our conclusions would likely have been different if we had only examined mean laying dates in relation to food and temperature and not examined the effects of wind speed and food across the entire reproductive period.

### The importance of food varies by reproductive stage

Although food abundance is often thought to be important to reproductive success, it is often not directly measured, nor analyzed across the entire reproductive cycle to determine when it is important (e.g., during egg production or offspring development). In our study, food abundance only had a significant effect on reproductive (fledging) success early in the season through its direct effect on laying date; there was no effect of food abundance on later stages (i.e., clutch size, hatching or fledging success). This was surprising given the emphasis in the literature on the temporal mismatch between the peaks of food abundance and nestling growth. A few other studies of Tree Swallows have also suggested that food abundance during the laying period is relatively more important to fledging success than food during the nestling period (e.g., Dunn and Hannon, 1989; Imlay et al., 2017; LeClair, 2012); however, there are also several studies that have found a strong positive effect of food abundance during the nestling period on fledging success (Twinning et al., 2018; Berzins et al., 2021; see also Martay et al., 2023 for another swallow species).

It is likely that these differing results depend on the seasonal timing of peak food abundance which can lead to different effects on the production of eggs (laying date or clutch size) or the ability to feed nestlings.

### Seasonal variation in food may also be important to fledging success

There is evidence from across North America that food abundance for swallows varies between populations in terms of overall abundance and timing. Some populations show steady increases in biomass over the season, but others show an increase in May and then a decline later in June (e.g., see Nematocera trends in Figure 5A of Dunn et al., 2022). In Wisconsin, food abundance generally increased over the season to a plateau in early June when eggs hatch (Figure 5A). Thus, food abundance was relatively high at hatching and remained so during the nestling growth period. Habitat can also interact with time of the season, as food abundance (Diptera biomass) was greater in areas with less intensive agriculture in southern Quebec, but this only occurred late in the season (Paquette et al., 2012). These seasonal and habitat differences could produce some of the differences we see in the timing of the importance of food abundance at different study sites, even if overall average food abundance is the same. This information could be important to identifying the causes of population decline, given that fledging success is one of the strongest demographic drivers of Tree Swallow populations (Cox et al., 2020; Weegman et al., 2017)

### Geographic variation in the timing of food abundance

Studies of Tree Swallows in locations with different levels of food support the hypothesis that the abundance and timing of food is important to reproductive success. For example, a long-term study of Tree Swallows near Port Rowan, Ontario, Canada found that at one site (Sewage lagoon) with relatively high food abundance early in the season, the number of young fledged was primarily influenced by food abundance during laying through its direct effects on clutch size and hatching success, rather than conditions during the nestling period (LeClair, 2012). However, at a nearby site (Long Point) with lower food abundance at the beginning of the season, temperature during the nestling period had the strongest effect on the number of young fledged (LeClair, 2012). Food abundance was relatively similar between these two sites during the nestling period and had no significant effects on the number of young fledged (LeClair, 2012). Male removal experiments in Tree Swallows also suggest that food abundance can be important to reproductive success. Removing male assistance with feeding nestlings had a stronger negative effect on fledging success in an area of lower than higher food abundance (Dunn & Robertson, 1992).

### Food availability is dependent on temperature and wind

Food abundance for swallows and other aerial insectivores is dependent on both temperature and wind. In terms of temperature, many climate change studies have focused on extreme weather events, such as heat waves and cold snaps, which can have detrimental effects on reproduction (van de Pol et al., 2017; Satarkar et al., 2026). In Tree Swallows, cold snaps or inclement weather (rainy and cold) during the early nestling period have been implicated in poor reproductive success in long-term studies in the northeastern United States (Blyth et al., 2019; Twining et al., 2018; Shipley et al., 2020) and southern Ontario (Cox et al., 2020) and Quebec (Garrett et al., 2022b). For example, in Quebec, fledging success was 36% lower when low temperatures (cold for 4 days) were combined with rainfall and more intensive agriculture in the landscape, all of which reduced food (Diptera) abundance (Garrett et al., 2022b). In contrast, we found no effect of the number of cold days on fledging success. The number of cold days did not appear to differ between Sherbrooke, QC and Saukville, WI as we found that the average female experienced 0.61 cold days (*±* 0.83 SD; 0.035 se, n= 548 complete females) when their nestlings were 1-12 days old compared with 0.9 *±* 1.1 days (SD) in Quebec (Garrett et al., 2022b). However, in contrast to the Ithaca, NY population, which has seen an increase in the number of cold days (Shipley et al., 2020), the Saukville, WI population has experienced a decrease during nestling days 1-6 (*b* = –0.079 *±*0.008 SE, *n* = 548 nests with complete data, *t* = 10.1, *p <* 0.001). Also, the proportion of complete nest failures has decreased from 19% in 2000 to zero in 2021 in Wisconsin (*b* = –0.009 *±* 0.00167, *t* = 5.63, *n* = 14 years, *p* <0.001, *r*^2^ = 0.73), compared with an increase in nest failures from 16 to 33% (1989 to 2015) in Ithaca, NY (Shipley et al., 2020). These results point to major differences between long-term study populations; understanding these differences will require combining data from several study sites and analyzing them within a standard framework.

The global decline in wind speed, and its ecological effects, has received much less attention than temperature change. Over land, the global decline in wind speed likely started in the 1960’s, although it has only been widely recognized by researchers for the last 20 years (Wu et al., 2025). The causes of the decline are generally attributed to reduced temperature gradients in the northern hemisphere caused by global warming (Karnauskas et al., 2018). Long-term atmospheric cycles such as the North Atlantic Oscillation, and increases in plant biomass and urbanization that increase surface roughness and, thus, slow wind speed, may also contribute to slowing wind speed (Wu et al., 2025). Although there have been many studies by climatologists and the wind energy industry, relatively few studies have examined the effects of slowing wind speed on the reproductive success of vertebrates. One example is a study of Tree Swallows in Alaska where wind speed has slowed by 21% over 16 years (2000-2015; Irons et al., 2017). This decline in wind speed was associated with earlier laying and hatching dates, while temperature had relatively little effect. Food abundance and other reproductive parameters such as clutch size and fledging success were not reported in this Alaskan study, but based on our results, it is likely that slower wind speed increases food availability and the rate of parental provisioning (Rose, 2009), which leads to greater fledging success (Taff et al., 2026b). Similarly, in a study of barn swallows (*Hirundo rustica*) in Denmark, there has been a long term decrease in wind speed in July, which is correlated with greater flying insect abundance and fledging success of second broods (Møller, 2013). Thus, changes in wind speed are widespread, and they are likely to be important to reproductive success in other populations and species of aerial insectivores.

### Why don’t populations change their timing of breeding?

Despite these changes in food abundance and weather, some populations do not appear to be changing their timing of breeding, even though there is apparent selection for breeding earlier. One explanation is that climate change allows late nesting birds to advance laying dates, but early nesting birds cannot advance as much because there is a constraint on laying earlier at the beginning of the season (due to poor weather or food conditions). As a consequence, the distribution of laying dates narrows (in the right hand tail), but the mean and earliest dates do not change significantly; hence the apparent stasis of laying date. Our study is consistent with this hypothesis as we found that only late nesting birds were advancing their laying dates, and there also appeared to be constraints on laying earlier than the first week of May (4 May was the earliest nest). Changepoint analyses revealed large increases in mean and minimum daily temperature on 29-30 Apr and food abundance showed increases on 3 and 20 May; interestingly the latter date is close to the long-term average laying date (18 May). It will be worth investigating whether these increases act as thresholds for birds to start laying in other populations as the laying date of Tree Swallows seems to show a similar response to temperature across their range (Taff et al., 2026a). Winkler et al. (2002) also found a similar narrowing of laying dates in relation to temperature in an analysis of Tree Swallows across North America, but in this case the mean laying date was advancing (not static as in this study) and they did not explore the causes of the constraint in detail. Many studies mention that laying earlier in the season might be constrained due to cold weather or lack of food, and several have shown that laying is more variable in warmer springs; however, few of these studies have actually measured how food and weather affect the distribution of laying (at the beginning and end). In the case of Tree Swallows most of the changes were at the right hand tail of the distribution with the late nesting birds, rather than the early birds which appeared to be more constrained.

### Conclusions

Our 25-year study demonstrates that climate change is not necessarily affecting populations through a simple, uniform shift in average timing, but rather through changes in the entire distribution of breeding caused by advances in the right-tail of the distribution and a piling up of early breeding individuals against constraints (poor weather and food supply) early in the season. By examining food and several weather variables in a comprehensive analysis, we found that early May rainfall, wind speed, and food availability have a strong effect on the shape of the laying date distribution. These effects on the shape of the distribution were much stronger than their effects on the mean laying date, probably because the early season constraints restrict movement of the mean. The dramatic 47% reduction in wind speed has increased food availability and allowed later nesting birds to advance their nesting, but they run into an early-season constraint of temperature and food that does not appear to relax until early May. These findings underscore that focusing strictly on shifts in phenological means overlooks critical changes in the distribution, and that accounting for changes in the shape of distributions can sometimes reveal cryptic effects that are important for accurately predicting the response of species to climate change.

## ACKNOWLEDGMENTS

We thank the UWM Field station and its staff, particularly G. Meyer, L. Nelson, J. Reinartz and R. Tagye, for logistical help over 26 years of field work. We also thank the many field assistants who helped collect nesting and insect abundance data. We thank D. Hussell for providing the tow nets and D. Winkler for the suction trap. Our research at the UWM Field Station was supported by grants from the College of Letters and Science, UWM and the Summer Undergraduate Research Fund. All procedures were conducted with IACUC approval from the University of Wisconsin Milwaukee and appropriate Federal and State permits.

## AUTHOR CONTRIBUTIONS

POD and LAW designed the study and collected the data. POD analyzed the data and wrote the first draft. Both authors edited and approved the final version.

## DATA AVAILABILITY STATEMENT

All data, including nest, weather and insect food abundance data, and reproducible workflows in R have been deposited in the Zenodo data repository (10.5281/zenodo.23676019) and will be made publicly available following publication of the manuscript.

