## Supplemental_Material for "A 47% decline in wind speed shapes phenology and selection on mismatching in Tree Swallows across a quarter century"

1372 **SUPPORTING INFORMATION** for review and publication

1376 **Appendix S1**

1378 **1. Imputing Nematoceran biomass for missing days of sampling**

Sampling with the suction trap occurred on an average of 65 days per season, and there
1380 were 956 days of sampling across all 15 years with data (2000-2021). There were 20
missing days of insect sampling in the first 4-6 days in May during 2000-2003, so we
1382 imputed values for Nematoceran biomass using predicted values from a regression
equation that included average rain during the 10 days prior to insect sampling, average
1384 wind speed (m/s) on the day of insect sampling and maximum temperature on the day of
insect sampling. The equation was: Nematoceran biomass (mg/day) = $1.576 + 10d \text{ rain}$
1386 $(0.289) + \text{mean wind speed} * -1.513 + \text{daily maximum temperature} * 0.238$. The equation
had a low predictive ability ($r^2 = 0.113$, $F_{3, 946} = 40.49$, $p < 0.001$), but excluding these imputed
1388 values did not have a qualitative effect on the results. We also examined a model with three
additional variables (julian date, solar radiation on the day of insect sampling and for the
1390 previous 10 days). Although all of these variables were significant, it only increased
predictive ability by 1.4% to 12.7%.

1392

### 2. Calculations for percent change in the mean and shape of the laying date distributions in Figures 5, 6.

#### General formula for calculating percent change for the 1.5 SD Shape Comparisons:

$$\%Change = (e^{\beta \times 1.5} - 1) \times 100\%$$

Beta values are from the full Bayesian gamma model in Table S5.

##### Figure 5A: Rainfall (beta = +0.729)

$$(e^{0.729 \times 1.5} - 1) \times 100 = (2.9847 - 1) \times 100 = +198\%$$

##### Figure 5B: Wind Speed (beta = -0.682)

$$(e^{-0.682 \times 1.5} - 1) \times 100 = (0.3595 - 1) \times 100 = -64\%$$

##### Figure 5C: Food Biomass (beta = -0.522)

$$(e^{-0.522 \times 1.5} - 1) \times 100 = (0.4570 - 1) \times 100 = -54\%$$

##### Figure 5D: Temperature (beta = -0.300)

$$(e^{-0.300 \times 1.5} - 1) \times 100 = (0.6376 - 1) \times 100 = -36\%$$

##### Figure 6A,B: food and wind interaction (beta = -0.714)

$$(e^{-0.714 \times 1.5} - 1) \times 100 = (0.3427 - 1) \times 100 = -66\%$$

### 3. Structural equation modeling

We used a structural equation model (SEM) to determine the relative importance of food and weather at various stages of reproduction. We constructed the SEM based on previous results and a priori hypotheses about the relationships between each environmental variable and reproduction. For consistency with the previous SEM study by LeClair (2012), we also used a similar SEM structure and variables from her analysis of two Tree Swallow populations near Long Point, Ontario, Canada. Methods were generally similar between studies but LeClair (2012) estimated food abundance using total insect biomass collected from tow-nets (Hussell & Quinney, 1985), while we used nematoceran biomass collected from suction traps. LeClair (2012) also had slightly different definitions of the reproductive periods than the ones we used above. The pre-laying period was the same in both LeClair

(2012) and this study (1-15 May), but the laying period was defined as 2 days before the 10<sup>th</sup> percentile of laying dates to the 90<sup>th</sup> percentile by LeClair (2012). For our main analyses, we followed our previous paper (Dunn et al., 2011) and used the 25<sup>th</sup> to 75<sup>th</sup> percentiles of laying dates, which was narrower and avoided a few days of overlap with the prelaying period. Incubation periods were defined the same in both studies (50<sup>th</sup> percentile of clutch completion dates to 50<sup>th</sup> percentile of hatch dates), while the nestling period was defined differently. The nestling period in LeClair (2012) was the 50<sup>th</sup> percentile of hatching to 16 days past that date, which would put half of the nests to the point of fledging since nestlings can fledge after day 16 of age. Our analysis used a nestling period from the 25<sup>th</sup> percentile of hatching dates to 15 days past the 75<sup>th</sup> percentile of hatching dates. Similar to LeClair (2012) we also simplified the analysis by only using known (banded) adult (ASY/AHY) females with data on all reproductive parameters ($n = 548$).

We started with an SEM that contained food (Nematocera biomass), mean temperature and wind speed variables that were likely to affect each stage of reproduction; specifically, laying date, clutch size, number of eggs hatched (N hatch), number of young reaching 12 d of age (D12) and number of young fledged (N Fledge). We averaged food, temperature and wind speed for the four reproductive periods defined above: pre-laying, laying, incubation and the nestling period.

The initial full model (Figure 11A; Table S11) contained effects of temperature both directly on the reproductive parameter (e.g., laying date or clutch size) and indirectly through food abundance. These relationships were supported by our earlier analyses of the direct effect of temperature on laying date (e.g., Appendix S1:Table S4, Figure 2A) and its potential indirect effect via food abundance. Similarly, wind speed had effects on both laying date (Figure 3C) and food abundance (Figure 6), but we do not know the relative importance of direct effects (e.g., on ability to forage or thermoregulate) versus indirect effects (e.g., reducing the availability of flying insects), so we included direct effects of wind speed on each reproductive variable as well as indirect effects via food abundance (see Figure 12A for paths). After running the full model, we sequentially removed non-significant paths until no non-significant paths remained (i.e., Figure 12B). Analyses were performed primarily in JMP with the Structural Equation Models procedure, because it provides more comprehensive graphing and statistical output, including automatic estimation of direct, indirect (Table S11) and total (Table S12) effects, which require further computations when using R packages, such as lavaan (Rosseel, 2012) and piecewiseSEM (Lefcheck, 2016). For reproducibility, we also analyze the same models with lavaan and piecewiseSEM using the R code in the Zenodo repository (10.5281/zenodo.23676019), and we present the results of the different methods in Table S14. Results were very similar as R-squared

1462 values between estimates from JMP and lavaan or pSEM were $>0.85$, while the r-squared
value for the comparison of estimates from lavaan and pSEM was $>0.99$ (Table S14).

1464

1466 **SUPPLEMENTARY FIGURES**

1468 **Figure S1.** Changepoints in mean and minimum temperature and food abundance during
1470 April to June. Panels show the daily temperature or Nematoceran biomass (dots) and the
1472 Red vertical dashed lines indicate the changepoint dates identified by the multiple change
points (mcp) package in R. Red bands indicate the confidence bands. Changepoints for
mean and minimum temperature were 29 Apr (119 Julian date) and 3 May (123 Julian date)
for Nematoceran biomass. There was a second changepoint for Nematoceran biomass at
20 May (140 Julian date).

1474

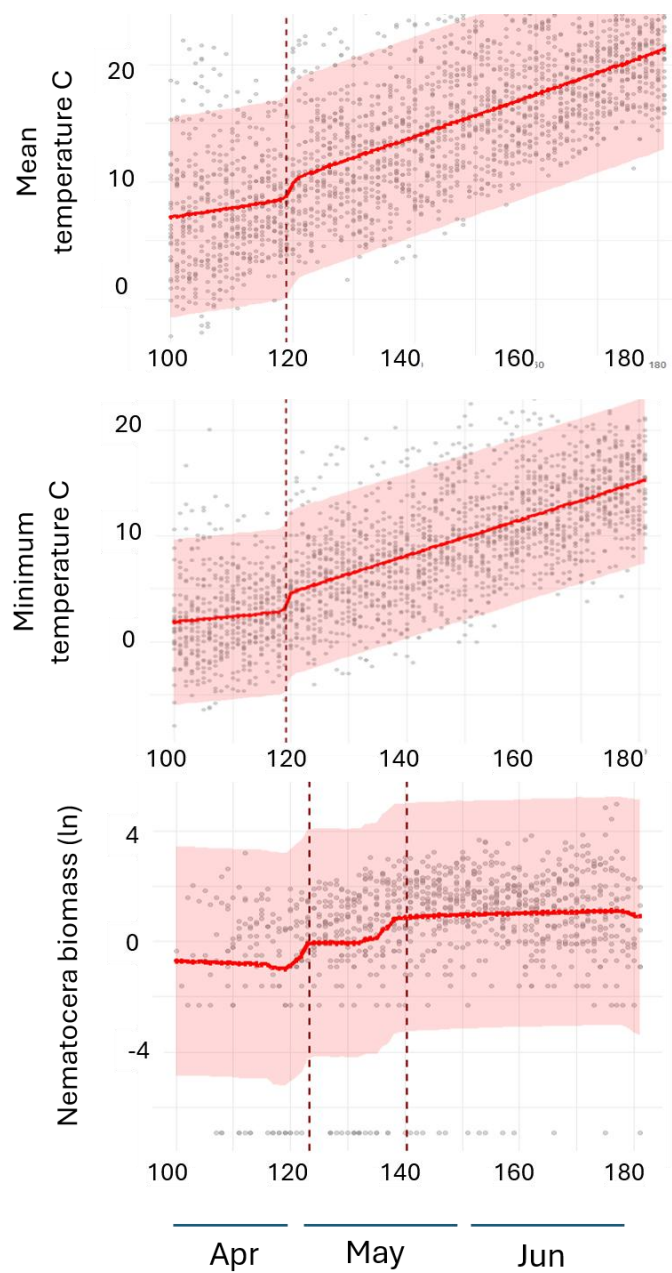

1476

1478 **Figure S2.** Change in the 5<sup>th</sup> percentile of egg laying dates from 2000 - 2021. The solid
1480 black line shows the estimated 5th percentile of egg-laying dates (Julian Day), estimated
1482 from the full quantile regression model (Table S7; year $p = 0.023$). The change becomes
1484 non-significant ($p = 0.378$) after dropping the interaction terms, which only increase
1486 predictive ability by 2.7% (Appendix S1: Table S7). In any case, the real-world change in
1488 laying date is only 1.8 days earlier across the 21 year period. To capture the change in wind
speed, the predictions integrate the observed annual mean pre-laying wind conditions
while holding all other weather and food variables constant at their long-term averages (0).
The shaded ribbon illustrates the 95% clustered bootstrap confidence interval (from 1,000
replications grouped by individual female id). The horizontal dark-red line at the bottom
indicates the earliest known date of egg laying (May 4<sup>th</sup>, Julian Day 124).

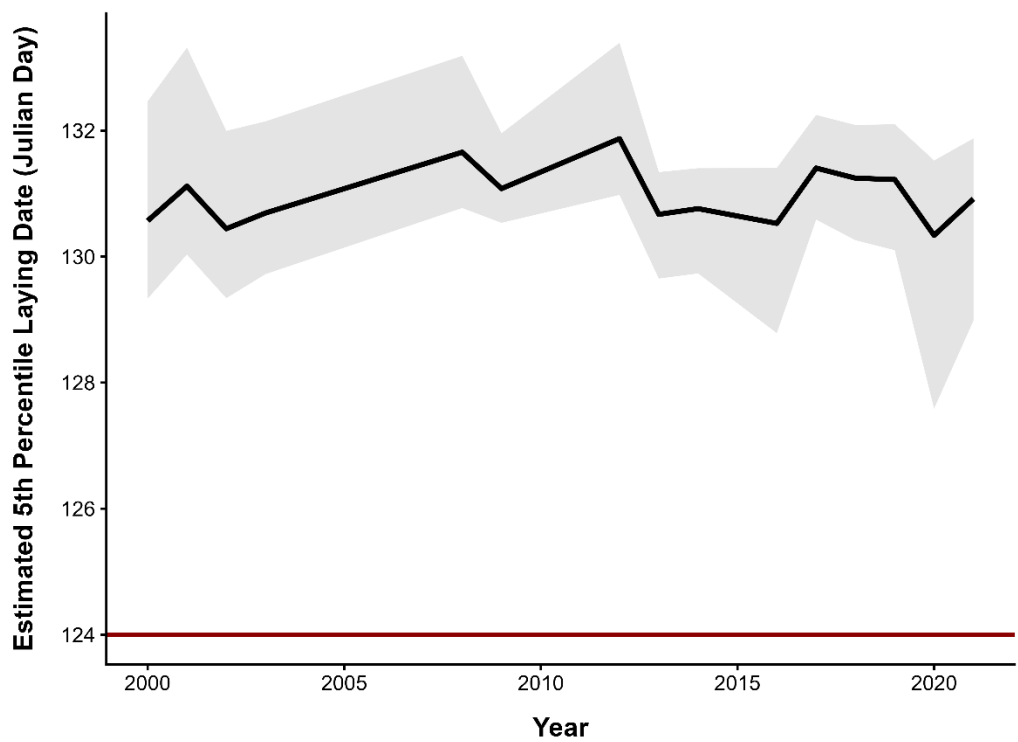

1490

1492

Table S1. Trends in annual laying and hatching dates (Y) versus year (X) N= 25. Laying dates are becoming less variable (lower SD) over time, because the laying dates of relatively late birds (75 - 95 perc) are advancing but the dates of early nesting birds are not shifting.

| Laying date parameter | slope | se | t | P | stand.<br>slope | r2 |
| --- | --- | --- | --- | --- | --- | --- |
| **All birds** |  |  |  |  |  |  |
| Mean | -0.144 | 0.088 | 1.64 | 0.114 | -0.324 | 0.105 |
| SD | -0.107 | 0.032 | -3.33 | 0.003 | -0.570 | 0.325 |
| Interquartile range (25-75 perc) | -0.201 | 0.060 | -3.38 | 0.003 | -0.576 | 0.332 |
| 5th percentile | -0.020 | 0.101 | -0.20 | 0.844 | -0.042 | 0.002 |
| 10th percentile | -0.016 | 0.096 | -0.16 | 0.872 | -0.034 | 0.001 |
| 25th percentile | -0.033 | 0.100 | -0.33 | 0.745 | -0.069 | 0.005 |
| **50th percentile** | -0.136 | 0.092 | 1.49 | 0.151 | -0.296 | 0.087 |
| 75th percentile | -0.245 | 0.094 | 2.59 | 0.016 | -0.476 | 0.226 |
| 90th percentile | -0.236 | 0.100 | 2.34 | 0.028 | -0.439 | 0.193 |
| 95th percentile | -0.407 | 0.100 | 4.06 | 0.001 | -0.646 | 0.417 |
| **ASY females only (n= 24 years)** |  |  |  |  |  |  |
| 50th percentile | -0.138 | 0.110 | 1.26 | 0.221 | -0.260 | 0.067 |
| 75th percentile | -0.300 | 0.107 | 2.81 | 0.010 | -0.513 | 0.263 |
| 90th percentile | -0.255 | 0.109 | 2.34 | 0.029 | 0.44664 | 0.199 |
| **Hatching date parameter (Y)** |  |  |  |  |  |  |
| Hatching date parameter (Y) | slope | se | t | P | stand.<br>slope | r2 |
| Mean | -0.130 | 0.089 | 1.450 | 0.160 | -0.290 | 0.084 |
| SD | -0.091 | 0.031 | -2.990 | 0.007 | -0.530 | 0.280 |
| 5th percentile | -0.006 | 0.100 | -0.060 | 0.953 | -0.012 | 0.001 |
| 10th percentile | -0.010 | 0.097 | -0.100 | 0.920 | -0.021 | 0.001 |
| 25th percentile | -0.059 | 0.097 | -0.61 | 0.548 | -0.126 | 0.016 |
| 50th percentile | -0.120 | 0.091 | 1.32 | 0.199 | -0.266 | 0.071 |
| 75th percentile | -0.256 | 0.083 | 3.07 | 0.005 | -0.539 | 0.291 |
| 90th percentile | -0.236 | 0.093 | 2.54 | 0.018 | -0.468 | 0.219 |

1496

1498

1500

Table S2. Annual change in weather and food during the pre-laying period (1-15 May) or on 18 May, the average laying date. Each weather or food variable was regressed on year using simple linear regressions (n = 26 for weather variables; N= 13 for food variables). Some variables were averaged for the 5 (5d ave) or 10 (10d ave) days before 18 May each year.

| Response variables (Y) | Estimate | Std<br>Error | t Ratio | Prob> \|t\| |
| --- | --- | --- | --- | --- |
| **Weather during pre-laying period** |  |  |  |  |
| Mean Temp 1-15 May | -0.065 | 0.050 | -1.3 | 0.2059 |
| Rain 1-15May | -0.071 | 0.052 | -1.36 | 0.1859 |
| Wind speed 1-15May | -0.049 | 0.006 | -7.9 | <.0001 |
| **Weather on the mean laying date (18 May)** |  |  |  |  |
| mean Temp DAILY | -0.044 | 0.106 | -0.41 | 0.6825 |
| Max Temp | -0.012 | 0.123 | -0.1 | 0.9209 |
| Min Temp | -0.077 | 0.098 | -0.79 | 0.4378 |
| total rain | -0.077 | 0.217 | -0.36 | 0.7252 |
| Mean WIND SPEED (m/s) | -0.045 | 0.021 | -2.13 | 0.0439 |
| mean Solar | -0.002 | 0.003 | -0.96 | 0.3452 |
| 5d prev ave temp | 0.077 | 0.079 | 0.98 | 0.3371 |
| 5d ave rain | -0.017 | 0.111 | -0.15 | 0.8783 |
| 5d ave wind | -0.035 | 0.011 | -3.15 | 0.0043 |
| 5d ave solar | -0.001 | 0.002 | -0.39 | 0.7012 |
| 10d prev ave temp | -0.004 | 0.062 | -0.07 | 0.9455 |
| 10d ave rain | -0.087 | 0.075 | -1.16 | 0.2561 |
| 10d ave solar | -0.002 | 0.002 | -0.9 | 0.3789 |
| 10d ave wind | -0.047 | 0.008 | -5.67 | <.0001 |
| Nematocera biomass | 0.094 | 0.151 | 0.62 | 0.5441 |
| 5d ave nemato | 0.052 | 0.132 | 0.39 | 0.7035 |

1502

1504

1506 Table S3. Annual change in weather and food on the long-term hatch date (6 June). Each
1508 weather or food variable (Y) was regressed on year (X) using simple linear regressions (n =
26 for weather variables; n = 13 for food variables). Some variables were averaged for the 5
(5d ave) or 10 (10d ave) day before 6 June each year.

| Response variable (Y) | Estimate | Std<br>Error | t Ratio | Prob> \|t\| |
| --- | --- | --- | --- | --- |
| mean Temp DAILY | 0.035 | 0.108 | 0.33 | 0.747 |
| Max Temp | 0.048 | 0.133 | 0.36 | 0.723 |
| Min Temp | 0.065 | 0.100 | 0.65 | 0.524 |
| total rain | 0.009 | 0.134 | 0.07 | 0.945 |
| Wind speed | -0.043 | 0.013 | -3.34 | 0.003 |
| mean Solar | 0.002 | 0.002 | 0.8 | 0.432 |
| 5d prev ave temp | 0.215 | 0.061 | 3.54 | 0.002 |
| 5d ave rain | -0.110 | 0.085 | -1.3 | 0.207 |
| 5d ave wind | -0.043 | 0.011 | -3.99 | 0.001 |
| 5d ave solar | 0.005 | 0.001 | 3.58 | 0.002 |
| 10d prev ave temp | 0.138 | 0.053 | 2.6 | 0.016 |
| 10d ave rain | -0.079 | 0.051 | -1.56 | 0.133 |
| 10d ave solar | 0.002 | 0.001 | 2.52 | 0.019 |
| 10d ave wind | -0.037 | 0.008 | -4.8 | <.0001 |
| Nematocera on day of laying | -0.174 | 0.351 | -0.5 | 0.628 |
| 5d ave Nematocera | 0.108 | 0.146 | 0.74 | 0.474 |

1510

1512 Table S4. Comparison of five Bayesian distributional models of laying date using Expected
1514 Log Predictive Density Difference (elpd), which measures the difference in predictive ability
1516 between models. The full model is given in Table S5 and includes parameters for the
location (mean) and shape (k) of the laying date distribution, weather and food variables for
1518 the prelaying period (absolute variables) and 5 days before the laying date of each female
(relative variables). The model without interactions did not contain the two interactions
between prelaying wind speed and year or food. The mean only model left out the shape
subcomponent of the full model, and the absolute only model left out the relative (5d
1520 previous) variables and only contained the prelaying variables that were the same for all
females in a given year (i.e., absolute). The timeline only model modeled year with a spline
1522 to capture non-linear trends; year was linear in the other models. All models contained a
code for each female (female\_id) as a random effect. All variables were standardized
1524 before analysis (using the “scale” function in R). Sample size was restricted to complete
cases with all weather and food data from 514 known females (N= 1029 nests over 15
1526 years). Models are arranged in increasing values of delta elpd from top to bottom.

| Model | elpd_diff | se_diff |
| --- | --- | --- |
| Full Distributional Model (Mean<br>& Shape) | 0 | 0 |
| Full_model no interactions | -76.2 | 14.5 |
| Traditional mean-only Model<br>(Constant Shape) | -215.7 | 27.4 |
| Full model no 5dbefore laying | -261.6 | 28.1 |
| Timeline only | -343.1 | 29.3 |

1528

1530 Table S5. The full Bayesian distribution models of laying date including parameters for the
1532 location (mean) and shape (k) of the laying date distribution, and weather and food
1534 variables for the prelaying period (absolute variables) and 5 days before the laying date of
1536 each female (relative variables). The model also included a random effect for each female
(n = 513 known females). Sample size was 1029 nests from 514 known females over 15
years, due to the smaller sample of years with insect data. All variables were standardized
before analysis (using the “scale” function in R). See Table S4 for a comparison of this
model with reduced models.

1538

| Predictor Variable | Posterior Mean | Lower 95% CI | Upper 95% CI | Est. Error (SD) | Interpretation of significant effects |
| --- | --- | --- | --- | --- | --- |
| Random Effect Hyperparameters (female_id) |  |  |  |  |  |
| sd(MEAN Intercept) | 0.044 | 0.021 | 0.070 | 0.012 | sign individual effects on mean |
| sd(SHAPE Intercept) | 1.197 | 0.966 | 1.452 | 0.122 | sign individual effects on shape |
| **MEAN** |  |  |  |  |  |
| Baseline Intercept | 2.769 | 2.741 | 2.796 | 0.014 |  |
| Nemato5dprev_scaled | 0.170 | 0.149 | 0.192 | 0.011 | Calendar effect: more food later in the season |
| Rain5dprev_scaled | -0.067 | -0.096 | -0.039 | 0.015 | More rain 5d before laying for early nesting birds |
| Temp5dprev_scaled | 0.089 | 0.072 | 0.106 | 0.009 | Calendar effect: warmer later in the season |
| Wind5dprev_scaled | 0.033 | 0.012 | 0.054 | 0.011 | More wind 5d before laying for late nesting birds |
| preLayNemato_scaled | -0.172 | -0.205 | -0.138 | 0.017 | More food prelaying associated with earlier laying |
| preLayRain_scaled | -0.024 | -0.054 | 0.005 | 0.015 |  |
| preLayTemp_scaled | -0.094 | -0.128 | -0.062 | 0.017 | Higher Temp prelaying = earlier laying |
| preLayWind_scaled | 0.179 | 0.120 | 0.238 | 0.030 | More wind prelaying = later laying |
| preLayWind_scaled : preLayNemato_scaled | 0.080 | 0.056 | 0.105 | 0.012 | High wind speed opposes the advancement due to food (preLayNemato = - 0.17) |
|  |  |  |  |  | see Figure 6D. Mean laying date is actually later in more recent years after accounting for environmental variables and shape difference |
| year_scaled | 0.065 | 0.003 | 0.126 | 0.032 |  |
| year_scaled :<br>preLayWind_scaled | 0.023 | -0.003 | 0.047 | 0.013 |  |
| **SHAPE** |  |  |  |  |  |
| Baseline Intercept | 3.123 | 2.834 | 3.425 | 0.150 |  |
| Nemato5dprev_scaled | 0.912 | 0.669 | 1.167 | 0.127 | More food before laying of individual leads to more synchronized laying |
| Rain5dprev_scaled | -0.369 | -0.505 | -0.228 | 0.072 |  |
| Temp5dprev_scaled | -0.361 | -0.514 | -0.212 | 0.077 |  |
| Wind5dprev_scaled | -0.188 | -0.413 | 0.034 | 0.114 |  |
| preLayNemato_scaled | -0.522 | -0.775 | -0.268 | 0.129 | More prelaying food leads to less synchronized laying |
| preLayRain_scaled | 0.729 | 0.523 | 0.934 | 0.104 | Rain leads to more synchronized laying |
| preLayTemp_scaled | -0.300 | -0.493 | -0.106 | 0.099 |  |
| preLayWind_scaled | -0.682 | -1.032 | -0.327 | 0.178 |  |
| preLayWind_scaled :<br>preLayNemato_scaled | -0.714 | -0.981 | -0.447 | 0.137 | Laying becomes more synchronized when food is abundant and wind speed is low.(Figure 6A, B) |
| year_scaled | -0.240 | -0.634 | 0.161 | 0.202 |  |
| year_scaled :<br>preLayWind_scaled | 0.291 | 0.086 | 0.502 | 0.108 | Long-term decline in wind speed is associated with compressed laying (Figure 6C, D) |

Table S6. Changepoint analysis of temperature and food (Nematocera) abundance.

Analyses used the multiple change points (mcp) package in R (Lindeløv 2020). Mcp allowed us to find changepoints along a day of the year timeline while incorporating year-to-year shifts by adding year as a random effect in the model. The mcp package does not explicitly test for the number of changepoints, so we compared models with 0, 1 and 2 changepoints using the loo package (Vehtari et al. 2017), which performs leave-one-out cross-validation and calculates Expected Log Predictive Density Difference (elpd), which measures the difference in predictive ability between Bayesian models. The top model received a elpd value of 0. Dates of breakpoints and their upper and lower 95% CI are given to the right.

|  |  | Number of changepoints |  |  |  |  |  |
| --- | --- | --- | --- | --- | --- | --- | --- |
| variable | elpd | 0 | 1 | 2 | date of breakpoint | lower | upper |
| mean temperature |  |  |  |  |  |  |  |
|  | elpd_diff | -5.6 | 0 | -4.8 | 119 | 113 | 123 |
|  | se_diff | 3.5 | 0 | 1.3 | 29-Apr | 22-Apr | 3-May |
| minimum temperature |  |  |  |  |  |  |  |
|  | elpd_diff | -8.1 | 0 | -4.3 | 119 | 118 | 121 |
|  | se_diff | 3.9 | 0 | 1.3 | 30-Apr | 29-Apr | 1-May |
| Nematocera |  |  |  |  |  |  |  |
|  | elpd_diff | -24.4 | -4.3 | 0 | 123 | 112.3 | 139.0 |
|  | se_diff | 7.5 | 5.9 | 0 | 3-May | 22-Apr | 19-May |
| second nematocera changepoint |  |  |  |  | 140 | 120 | 180 |
|  |  |  |  |  | 20-May |  |  |

1554

1556 Table S7. Analysis of the 5<sup>th</sup> percentile of laying dates using quantile regression (quantreg
1558 package in R). Julian laying date was the response variable and all predictors were mean-
1560 centered and scaled in R. The 5<sup>th</sup> percentile advanced by 1.8 days over the 21 year period
(2000-2021). Appendix S1: Figure S2 shows the long-term trend in the 5<sup>th</sup> percentile based
1562 on the full model below. The variables in the full model with interactions were chosen
based on the model in Appendix S1: Table S5. Although the full model is significantly better
than the main effects model (without interactions) based on a nested Wald test $F_{2, 1021} =$
18.0, $p < 0.001$), the gain in predictive ability was just 2.7% (pseudo-$R^2 = 0.027$).

1564

| Model and Parameter Group | Estimate ($\beta$) | se | t | p (>\|t\|) |
| --- | --- | --- | --- | --- |
| FULL MODEL WITH INTERACTIONS |  |  |  |  |
| Intercept | 130.6 | 0.326 | 400.4 | 0.000 |
| year_scaled | -1.28 | 0.564 | -2.27 | 0.023 |
| preLayTemp_scaled | -1.91 | 0.268 | -7.14 | 0.000 |
| preLayRain_scaled | -0.60 | 0.300 | -2.01 | 0.045 |
| preLayWind_scaled | -1.07 | 0.519 | -2.06 | 0.040 |
| preLayNemato_scaled | -1.29 | 0.337 | -3.83 | 0.000 |
| preLayWind_scaled:<br>preLayNemato_scaled | 1.38 | 0.287 | 4.81 | 0.000 |
| year_scaled:<br>preLayWind_scaled | -1.10 | 0.271 | -4.07 | 0.000 |
| MAIN-EFFECTS MODEL - NO INTERACTIONS |  |  |  |  |
| Intercept | 130.8 | 0.273 | 479.9 | 0.000 |
| year_scaled | -0.68 | 0.766 | -0.88 | 0.378 |
| preLayTemp_scaled | -0.97 | 0.205 | -4.75 | 0.000 |
| preLayRain_scaled | -0.02 | 0.289 | -0.08 | 0.936 |
| preLayWind_scaled | -0.81 | 0.738 | -1.10 | 0.270 |
| preLayNemato_scaled | -0.50 | 0.359 | -1.39 | 0.166 |

1566

1568

1570

1572

Table S8. Best predictors of the peak date of Nematoceran biomass (annual estimates) from a sliding window analysis of weather variables using the climwin package in R. Climwin examined windows up to 84 days before 23 June (ie, up to ~1 Apr) with a 2 day min window.

| Weather variable | time window | R<sup>2</sup> | slope | se | deltaAICc | raw *p* |
| --- | --- | --- | --- | --- | --- | --- |
| MaxT | 66 - 54 (18- 30 Apr) | 0.624 | 6.2 | 1.33 | -11.49 | <0.001 |
| Min T | 60 - 57 (24-27 Apr) | 0.567 | 5.5 | 1.33 | -9.4 | 0.001 |
| Wind | 58 - 54 (26- 30 Apr) | 0.392 | 23.4 | 8.09 | -4.3 | 0.013 |
| Rain | 84 - 58 (1- 26 Apr) | 0.632 | 8.0 | 1.68 | -11.8 | <0.001 |
| Solar | 28 - 1 (26 May - 22 Jun) | 0.329 | 147.7 | 58.48 | -2.8 | 0.025 |

1574

1576

1578

1580

Table S9. Selection gradient analysis of phenological mismatch (the difference between dates of hatch and peak food abundance) over a 22-year time span. Results are from a Bayesian distributional model using brms. Data are from 932 nests and 15 years.

| Predictors | Relative Fledging Success (w) |  |  |
| --- | --- | --- | --- |
|  | Estimates | SE | CrI (95%) |
| Baseline Intercept | 1.12 | 0.05 | 1.02 – 1.24 |
| (Adult ASY Females) |  |  |  |
| Mismatch (Directional) | 0.01 | 0.04 | -0.07 – 0.09 |
| Mismatch<sup>2</sup> (Stabilizing) | **-0.08** | 0.04 | -0.17 – -0.00 |
| Decade (Year/10) | -0.06 | 0.04 | -0.14 – 0.01 |
| Female Age Class (SY Younger) | -0.11 | 0.06 | -0.23 – 0.00 |
| Female Age Class (Unknown) | **-0.10** | 0.04 | -0.18 – -0.02 |
| Mismatch: Decade | 0.01 | 0.03 | -0.06 – 0.07 |
| (Directional Change) |  |  |  |
| Mismatch<sup>2</sup>: Decade | **0.06** | 0.03 | 0.00 – 0.13 |
| (Stabilizing change) |  |  |  |

1582

1584

Table S10. Selection gradient analysis of laying date over a 22-year time span. Results are from a Bayesian distributional model using brms. Data are from 1351 nests and 22 years.

| Predictors | Relative Fledging Success (w) |  |  |
| --- | --- | --- | --- |
|  | Estimates | SE | CrI (95%) |
| Intercept (Adult ASY Females) | **1.06** | 0.03 | 0.99 – 1.12 |
| Laying Date (Directional) | 0.06 | 0.03 | -0.01 – 0.13 |
| Laying Date<sup>2</sup> (Stabilizing) | -0.03 | 0.02 | -0.07 – 0.00 |
| Decade (Year/10) | -0.02 | 0.02 | -0.07 – 0.02 |
| Female Age Class (SY Younger) | -0.11 | 0.07 | -0.24 – 0.02 |
| Female Age Class (Unknown) | -0.05 | 0.03 | -0.10 – 0.00 |
| Laying Date:Decade<br>(Directional Change) | -0.03 | 0.03 | -0.09 – 0.02 |
| Laying Date<sup>2</sup>:Decade<br>(Stabilizing change) | 0.02 | 0.02 | -0.01 – 0.05 |

1586

Table S11. Statistical details of SEM analyses of Tree Swallow fledging success (n Fledged) shown in Figure 12. Estimates are standardized values and are sorted by outcome and then $p$ value (bold for $p < 0.05$). Analyses were performed in JMP 19 (JMP 19 2026). The full model in Fig. 12A had an AICc = 23,480, CFI = 0.271 and RMSEA = 0.384. The final reduced model in Fig. 12B had an AICc = 13,781, CFI = 0.707 and RMSEA = 0.203. R-squared values for both models were similar: 0.15 for laying date(LayDate), 0.04 for clutch size (CS), 0.35 for number of hatched eggs(Hatch), 0.51 for number of nestlings on day 12 (Nd12), 0.86 for number of young fledged (Fledge) and 0.07 for prelaying Nematocera biomass(Nemat1\_15May). Variables names contain 1\_15May for the pre-laying period, Lay for the laying period, Incub for the incubation period and Nestl for the nestling period. Definitions of the time periods follow LeClair (2012) and are described in Appendix S1. See Table S12 for the total effects.

|  |  | Full model (Fig. 11A) |  |  | Reduced model (Fig. 11B) |  |  |
| --- | --- | --- | --- | --- | --- | --- | --- |
| | | Prob>\|Z\| | | | | | |
| predictor | outcome | full | Estimate | SE | Prob>\|Z\| | Estimate | SE |
| Direct paths |  |  |  |  |  |  |  |
| sLayDate | sCS | 0 | -0.157 | 0.043 | 0.0001 | -0.155 | 0.042 |
| sLayWind | sCS | 0.001 | -0.135 | 0.042 | 0.001 | -0.136 | 0.041 |
| sLayNemato | sCS | 0.572 | 0.024 | 0.042 |  |  |  |
| sLayT | sCS | 0.885 | -0.006 | 0.044 |  |  |  |
| sNd12 | sFledge | 0 | 0.929 | 0.006 | 0.0001 | 0.927 | 0.006 |
| sNestlT | sFledge | 0.002 | -0.05 | 0.016 |  |  |  |
| sNestl_Nemato | sFledge | 0.544 | -0.011 | 0.019 |  |  |  |
| sNestlWind | sFledge | 0.933 | -0.001 | 0.016 |  |  |  |
| sCS | sHatch | 0 | 0.572 | 0.029 | 0.0001 | 0.576 | 0.029 |
| sIncubT | sHatch | 0 | 0.145 | 0.039 | 0.0001 | 0.139 | 0.035 |
| sIncub_Nemato | sHatch | 0.137 | 0.06 | 0.04 |  |  |  |
| sIncubWind | sHatch | 0.774 | 0.012 | 0.04 |  |  |  |
| sIncubT | sIncub_Nemato | 0 | 0.3 | 0.018 |  |  |  |
| sIncubWind | sIncub_Nemato | 0 | -0.181 | 0.028 |  |  |  |
| sLN_Nemat1_15May | sLayDate | 0 | -0.25 | 0.04 | 0.0001 | -0.25 | 0.04 |
| sTemp1_(1-15May) | sLayDate | 0 | -0.312 | 0.04 | 0.0001 | -0.312 | 0.04 |
| sWIND_1-15May | sLayDate | 0.013 | 0.103 | 0.042 | 0.0129 | 0.103 | 0.042 |
| sLayT | sLayNemato | 0 | -0.158 | 0.041 |  |  |  |
| sLayWind | sLayNemato | 0.026 | -0.093 | 0.042 |  |  |  |
| sTemp1_(1-15May) | sLN_Nemat1_15May | 0 | -0.184 | 0.043 | 0.0001 | -0.184 | 0.043 |
| sWIND_1-15May | sLN_Nemat1_15May | 0 | -0.192 | 0.043 | 0.0001 | -0.192 | 0.043 |
| sHatch | sNd12 | 0 | 0.717 | 0.021 | 0.0001 | 0.72 | 0.02 |
| sNestlWind | sNd12 | 0.019 | -0.071 | 0.03 |  |  |  |

|  |  |  |  |  |
| --- | --- | --- | --- | --- |
| sNestlT | sNd12 | 0.122 | 0.046 | 0.03 |
| sNestl_Nemato | sNd12 | 0.409 | -0.028 | 0.034 |
| sNestlT | sIncub_Nemato | 0 | 0.288 | 0.016 |
| sNestlWind | sIncub_Nemato | 0 | -0.193 | 0.026 |

##### Indirect effects

| Predictor | outcome | Prob>\|Z\|<br>full | Estimate | SE | Prob>\|Z\| | Estimate | SE |
| --- | --- | --- | --- | --- | --- | --- | --- |
| sLN_Nemat1_15May | sCS | 0.002 | 0.04 | 0.013 | 0.002 | 0.039 | 0.012 |
| sTemp1_(1-15May) | sCS | 0.002 | 0.04 | 0.013 | 0.002 | 0.039 | 0.012 |
| sWIND_1-15May | sCS | 0.012 | -0.024 | 0.009 | 0.011 | -0.024 | 0.009 |
| sLayT | sCS | 0.576 | -0.004 | 0.007 |  |  |  |
| sLayWind | sCS | 0.584 | -0.002 | 0.004 |  |  |  |
| sHatch | sFledge | 0 | 0.662 | 0.03 | 0 | 0.669 | 0.03 |
| sCS | sFledge | 0 | 0.377 | 0.028 | 0 | 0.383 | 0.029 |
| sIncubT | sFledge | 0 | 0.109 | 0.029 | 0 | 0.094 | 0.024 |
| sLayDate | sFledge | 0.001 | -0.059 | 0.017 | 0 | -0.059 | 0.017 |
| sLayWind | sFledge | 0.002 | -0.052 | 0.016 | 0.002 | -0.052 | 0.017 |
| sLN_Nemat1_15May | sFledge | 0.003 | 0.015 | 0.005 | 0.002 | 0.015 | 0.005 |
| sTemp1_(1-15May) | sFledge | 0.003 | 0.015 | 0.005 | 0.002 | 0.015 | 0.005 |
| sWIND_1-15May | sFledge | 0.013 | -0.009 | 0.004 | 0.013 | -0.009 | 0.004 |
| sNestlWind | sFledge | 0.036 | -0.058 | 0.028 |  |  |  |
| sIncub_Nemato | sFledge | 0.14 | 0.034 | 0.023 |  |  |  |
| sNestlT | sFledge | 0.294 | 0.033 | 0.031 |  |  |  |
| sNestl_Nemato | sFledge | 0.408 | -0.022 | 0.027 |  |  |  |
| sLayNemato | sFledge | 0.572 | 0.009 | 0.016 |  |  |  |
| sLayT | sFledge | 0.814 | -0.004 | 0.016 |  |  |  |
| sIncubWind | sFledge | 0.985 | 0 | 0.026 |  |  |  |
| sLayDate | sHatch | 0 | -0.089 | 0.025 | 0 | -0.088 | 0.025 |
| sLayWind | sHatch | 0.001 | -0.078 | 0.024 | 0.001 | -0.078 | 0.024 |
| sTemp1_(1-15May) | sHatch | 0.002 | 0.023 | 0.007 | 0.002 | 0.022 | 0.007 |
| sLN_Nemat1_15May | sHatch | 0.002 | 0.023 | 0.007 | 0.002 | 0.022 | 0.007 |
| sWIND_1-15May | sHatch | 0.013 | -0.014 | 0.005 | 0.012 | -0.013 | 0.005 |
| sIncubT | sHatch | 0.14 | 0.018 | 0.012 |  |  |  |
| sIncubWind | sHatch | 0.149 | -0.011 | 0.007 |  |  |  |
| sLayNemato | sHatch | 0.572 | 0.014 | 0.024 |  |  |  |
| sLayT | sHatch | 0.814 | -0.006 | 0.024 |  |  |  |
| sWIND_1-15May | sLayDate | 0 | 0.049 | 0.014 | 0 | 0.049 | 0.014 |
| sTemp1_(1-15May) | sLayDate | 0.001 | 0.044 | 0.013 | 0.001 | 0.044 | 0.013 |
| sCS | sNd12 | 0 | 0.406 | 0.03 | 0 | 0.413 | 0.03 |
| sIncubT | sNd12 | 0 | 0.117 | 0.031 | 0 | 0.101 | 0.025 |
| sLayDate | sNd12 | 0.001 | -0.063 | 0.018 | 0 | -0.063 | 0.018 |
| sLayWind | sNd12 | 0.002 | -0.056 | 0.018 | 0.002 | -0.056 | 0.018 |
| sTemp1_(1-15May) | sNd12 | 0.002 | 0.016 | 0.005 | 0.002 | 0.016 | 0.005 |
| sLN_Nemat1_15May | sNd12 | 0.003 | 0.016 | 0.005 | 0.002 | 0.016 | 0.005 |
| sWIND_1-15May | sNd12 | 0.013 | -0.01 | 0.004 | 0.013 | -0.01 | 0.004 |
| sIncub_Nemato | sNd12 | 0.14 | 0.036 | 0.024 |  |  |  |
| sNestlT | sNd12 | 0.408 | -0.009 | 0.01 |  |  |  |
| sNestlWind | sNd12 | 0.411 | 0.005 | 0.007 |  |  |  |
| sLayNemato | sNd12 | 0.572 | 0.01 | 0.017 |  |  |  |
| sLayT | sNd12 | 0.814 | -0.004 | 0.017 |  |  |  |
| sIncubWind | sNd12 | 0.985 | 0.001 | 0.028 |  |  |  |

1600

1602

1604

Table S12. Total effects from final (reduced) SEM analysis of Tree Swallow fledging success (N Fledged) shown in Figure 12B and Table S11. Estimates are standardized values and are sorted by outcome and then *p* value (bold for *p* < 0.05). Analyses were performed in JMP 19 (JMP 19 2026). Variables names have an “s” before each name to indicate that they are standardized and contain 1\_15May for the pre-laying period, Lay for the laying period, Incub for the incubation period and Nestl for the nestling period. Definitions of the time periods mostly follow LeClair (2012) and are described in the main text. See Table S11 for more details.

| Predictor | Outcome | Prob>\|Z\| | Estimate | Std Error | Wald Z |
| --- | --- | --- | --- | --- | --- |
| sLayDate | sCS | **0.000** | -0.153 | 0.042 | -3.656 |
| sLayWind | sCS | **0.001** | -0.136 | 0.042 | -3.250 |
| sTemp1_(1-15May) | sCS | **0.002** | 0.039 | 0.012 | 3.150 |
| sLN_Nemat1_15May | sCS | **0.002** | 0.039 | 0.012 | 3.139 |
| sWIND_1-15May | sCS | **0.011** | -0.024 | 0.009 | -2.541 |
| sCS | sFledge | **0.000** | 0.383 | 0.029 | 13.395 |
| sHatch | sFledge | **0.000** | 0.669 | 0.030 | 22.491 |
| sNd12 | sFledge | **0.000** | 0.928 | 0.016 | 58.122 |
| sIncubT | sFledge | **0.000** | 0.094 | 0.024 | 3.958 |
| sLayDate | sFledge | **0.000** | -0.059 | 0.017 | -3.527 |
| sLayWind | sFledge | **0.002** | -0.052 | 0.017 | -3.158 |
| sTemp1_(1-15May) | sFledge | **0.002** | 0.015 | 0.005 | 3.067 |
| sLN_Nemat1_15May | sFledge | **0.002** | 0.015 | 0.005 | 3.056 |
| sWIND_1-15May | sFledge | **0.013** | -0.009 | 0.004 | -2.497 |
| sCS | sHatch | **0.000** | 0.572 | 0.034 | 16.676 |
| sIncubT | sHatch | **0.000** | 0.140 | 0.035 | 4.020 |
| sLayDate | sHatch | **0.000** | -0.088 | 0.025 | -3.571 |
| sLayWind | sHatch | **0.001** | -0.078 | 0.024 | -3.190 |
| sTemp1_(1-15May) | sHatch | **0.002** | 0.022 | 0.007 | 3.095 |
| sLN_Nemat1_15May | sHatch | **0.002** | 0.022 | 0.007 | 3.084 |
| sWIND_1-15May | sHatch | **0.012** | -0.013 | 0.005 | -2.512 |
| sTemp1_(1-15May) | sLayDate | **0.000** | -0.254 | 0.041 | -6.209 |
| sLN_Nemat1_15May | sLayDate | **0.000** | -0.256 | 0.042 | -6.122 |
| sWIND_1-15May | sLayDate | **0.000** | 0.153 | 0.043 | 3.535 |
| sTemp1_(1-15May) | sLN_Nemat1_15May | **0.000** | -0.172 | 0.041 | -4.234 |
| sWIND_1-15May | sLN_Nemat1_15May | **0.000** | -0.190 | 0.043 | -4.429 |
| sCS | sNd12 | **0.000** | 0.413 | 0.030 | 13.766 |
| sHatch | sNd12 | **0.000** | 0.721 | 0.030 | 24.391 |
| sIncubT | sNd12 | **0.000** | 0.101 | 0.025 | 3.967 |
| sLayDate | sNd12 | **0.000** | -0.063 | 0.018 | -3.533 |
| sLayWind | sNd12 | **0.002** | -0.056 | 0.018 | -3.163 |
| sTemp1_(1-15May) | sNd12 | **0.002** | 0.016 | 0.005 | 3.071 |
| sLN_Nemat1_15May | sNd12 | **0.002** | 0.016 | 0.005 | 3.060 |
| sWIND_1-15May | sNd12 | **0.013** | -0.010 | 0.004 | -2.499 |

1614

1616

1618

1620 Table S13. Nesting success (binomial; 1=fledged at
1622 least one nesting) in relation to weather and food
1624 variables during incubation (Incub) and nestling (Nestl)
periods. $n = 548$ nests with complete reproductive
success data. FDR indicates $p$ values adjusted for the
False Discovery Rate. LN Nemato indicates log
transformed Nematoceran biomass.

1626

| Predictor | $p$ | FDR $p$ Value |
| --- | --- | --- |
| Mean Temp Incub period | 0.000 | 0.000 |
| Mean Wind Incub | 0.015 | 0.083 |
| Mean Wind Nestl | 0.023 | 0.083 |
| LN Nemato Incub Period | 0.080 | 0.220 |
| Mean Temp Nestl period | 0.348 | 0.547 |
| Mean Rain Incub | 0.252 | 0.547 |
| Nematocera Nestl period | 0.312 | 0.547 |
| Mean Rain Nestl | 0.631 | 0.771 |
| Mean Solar Incub | 0.574 | 0.771 |
| Mean Solar Nestl | 0.788 | 0.788 |
| LN Nemato Nestl Period | 0.784 | 0.788 |

1628

1630 Table S14. Comparison of results of SEM in Fig. 12A using JMP, lavaan and pSEM. $R^2$ values
1632 between estimates from JMP and lavaan or pSEM are $>0.85$, while $r^2$ values for lavaan and
pSEM are $>0.99$.

| Predictor | Outcome | JMP<br>Prob>\|Z\| | JMP<br>Estimate | lavaan<br>P(>\|z\|) | lavaan<br>Estimate | pSEM<br>P.Value | pSEM<br>Std.Estimate |
| --- | --- | --- | --- | --- | --- | --- | --- |
| sCS | sHatch | 0.000 | 0.572 | 0.000 | 0.570 | 0.000 | 0.573 |
| sHatch | sNd12 | 0.000 | 0.717 | 0.000 | 0.712 | 0.000 | 0.712 |
| sIncub_Nemato | sHatch | 0.137 | 0.060 | 0.160 | 0.051 | 0.151 | 0.051 |
| sIncubT | sHatch | 0.000 | 0.145 | 0.008 | 0.146 | 0.000 | 0.142 |
| sIncubT | sIncub_Nemato | 0.000 | 0.300 | 0.001 | -0.158 | 0.002 | -0.142 |
| sIncubWind | sHatch | 0.774 | 0.012 | 0.799 | 0.012 | 0.821 | 0.009 |
| sIncubWind | sIncub_Nemato | 0.000 | -0.181 | 0.000 | -0.293 | 0.000 | -0.266 |
| sLayDate | sCS | 0.000 | -0.157 | 0.000 | -0.156 | 0.000 | -0.167 |
| sLayNemato | sCS | 0.572 | 0.024 | 0.567 | 0.024 | 0.460 | 0.030 |
| sLayT | sCS | 0.885 | -0.006 | 0.888 | -0.006 | 0.762 | -0.013 |
| sLayT | sLayNemato | 0.000 | -0.158 | 0.000 | -0.156 | 0.000 | -0.159 |
| sLayWind | sCS | 0.001 | -0.135 | 0.001 | -0.135 | 0.006 | -0.113 |
| sLayWind | sLayNemato | 0.026 | -0.093 | 0.074 | -0.093 | 0.028 | -0.093 |
| sLN_Nemat1_15May | sLayDate | 0.000 | -0.250 | 0.000 | -0.256 | 0.000 | -0.269 |
| sNd12 | sFledge | 0.000 | 0.929 | 0.000 | 0.929 | 0.000 | 0.929 |
| sNestl_Nemato | sFledge | 0.544 | -0.011 | 0.531 | -0.009 | 0.546 | -0.010 |
| sNestl_Nemato | sNd12 | 0.409 | -0.028 | 0.456 | -0.024 | 0.410 | -0.025 |
| sNestlT | sFledge | 0.002 | -0.050 | 0.002 | -0.052 | 0.002 | -0.050 |
| sNestlT | sNd12 | 0.122 | 0.046 | 0.134 | 0.048 | 0.124 | 0.046 |
| sNestlT | sNestl_Nemato | 0.000 | 0.288 | 0.000 | -0.097 | 0.229 | -0.050 |
| sNestlWind | sFledge | 0.933 | -0.001 | 0.901 | -0.001 | 0.933 | -0.001 |
| sNestlWind | sNd12 | 0.019 | -0.071 | 0.058 | -0.071 | 0.019 | -0.071 |
| sNestlWind | sNestl_Nemato | 0.000 | -0.193 | 0.000 | -0.144 | 0.002 | -0.131 |
| sTemp1_(1-15May) | sLayDate | 0.000 | -0.312 | 0.000 | -0.298 | 0.000 | -0.314 |
| sTemp1_(1-15May) | sLN_Nemat1_15May | 0.000 | -0.184 | 0.000 | -0.172 | 0.000 | -0.185 |
| sWIND_1-15May | sLayDate | 0.013 | 0.103 | 0.022 | 0.105 | 0.019 | 0.107 |
| sWIND_1-15May | sLN_Nemat1_15May | 0.000 | -0.192 | 0.000 | -0.190 | 0.000 | -0.198 |

1638

1656

1658
